# The luminal AAA+ ATPase torsinA mediates distinct mechanisms of nuclear-cytoplasmic communication by adopting different functional assembly states

**DOI:** 10.1101/2021.12.09.472013

**Authors:** Kwang-Ho Hur, Jared Hennen, Cosmo A. Saunders, Amy Schoenhoefen, Patrick T. Willey, Cassandra M. Morris, Rabab H. Ibrahim, Ravi S. Maisuria, Joachim D. Mueller, G.W. Gant Luxton

## Abstract

Chemical and mechanical nuclear-cytoplasmic communication across the nuclear envelope (NE) is largely mediated by the nuclear pore complex (NPC) and the linker of nucleoskeleton and cytoskeleton (LINC) complex, respectively. While NPC and LINC complex assembly are functionally related, the mechanisms responsible for this relationship remain poorly understood. Here, we investigated how the luminal ATPases associated with various cellular activities (AAA+) protein torsinA promotes NPC and LINC complex assembly using fluorescence fluctuation spectroscopy (FFS), quantitative photobleaching analyses, and functional cellular assays. We report that torsinA controls LINC complex-dependent nuclear-cytoskeletal coupling as a soluble hexameric AAA+ protein and interphase NPC biogenesis as a membrane-associated helical polymer. These findings help resolve the conflicting models of torsinA function that were recently proposed based on *in vitro* structural studies. Our results will enable future studies of the role of defective nuclear-cytoplasmic communication in DYT1 dystonia and other diseases caused by mutations in torsinA.

## INTRODUCTION

The nuclear envelope (NE) is a specialized subdomain of the endoplasmic reticulum (ER) that surrounds the genome and defines the nuclear compartment in eukaryotic cells (Kite, 1913). It is composed of concentric inner and outer nuclear membranes (INM and ONM, respectively), which are separated by a perinuclear space that is contiguous with the ER lumen (Watson, 1955). The INM possesses a distinct set of proteins that interact with the underlying nuclear lamina and chromatin, whereas the ONM is an extension of the ER (Burke and Stewart, 2014). Fusion between the INM and the ONM creates fenestrations through the NE that are occupied by nuclear pore complexes (NPCs) (Watson, 1959).

Bidirectional nuclear-cytoplasmic communication, which can be either biochemical or mechanical in nature, is critical for cellular function and homeostasis (Knockenhauer and Schwartz, 2016). The chief mediators of nuclear-cytoplasmic communication are NPCs and linker of nucleoskeleton and cytoskeleton (LINC) complexes. Both NPCs and LINC complexes are multiprotein NE-spanning molecular bridges (Jahed et al., 2016). NPCs are the largest macromolecular complexes present in the NE, being made up of ∼30 different types of proteins known as nucleoporins (Fernandez-Martinez and Rout, 2021). In contrast, LINC complexes are composed of the ONM Klarsicht/ANC-1/SYNE homology (KASH) proteins and the INM Sad1/UNC-84 (SUN) proteins, which directly interact within the perinuclear space of the NE (Starr and Fridolfsson, 2010). Interestingly, LINC complex proteins are required for NPC assembly and their even distribution throughout the NE as well as their ability to perform nucleo-cytoplasmic transport (Liu et al., 2007; Talamas and Hetzer, 2011; Li and Noegel, 2015). In addition, LINC complex assembly requires the successful targeting of SUN proteins to the INM via their transport through NPCs (Tapley et al., 2011; Zuleger et al., 2011; Ungricht et al., 2015). Moreover, both NPCs and LINC complexes are necessary for proper NE architecture (Crisp et al., 2006; Cain et al., 2014; Zhou and Panté, 2010; Patel and Rexach, 2008; Vollmer et al., 2015; Mészáros et al., 2015). Despite these intriguing findings and the importance of NPCs and LINC complexes for cellular physiology, the mechanisms underlying the functional relationship between these NE-spanning molecular bridges remain unknown. To better understand the functional relationship between LINC complexes and NPCs, we sought to determine how the luminal ATPases associated with various cellular activities (AAA+) protein torsinA promotes their assembly from within the perinuclear space of the NE.

TorsinA was first identified in 1997 as the protein encoded by the causative gene of the debilitating and currently incurable neurological movement disorder early-onset primary (DYT1) dystonia (Ozelius et al., 1997). TorsinA is a conserved essential protein that has been implicated in both LINC complex and NPC assembly (Rampello et al., 2020b). More specifically, torsinA interacts with KASH and SUN proteins within the perinuclear space (Chalfant et al., 2019; Nery et al., 2008; Huelgas-Morales et al., 2020) and is required for the physical coupling of the nucleus and the cytoskeleton (Saunders et al., 2017). In addition, torsinA is required for the biogenesis of functional NPCs in interphase cells (Laudermilch et al., 2016; Pappas et al., 2018; Rampello et al., 2020a; VanGompel et al., 2015; Vander Heyden et al., 2009; Dominguez Gonzalez et al., 2018). AAA+ proteins typically function as higher-order oligomeric (e.g. hexameric rings or double rings) molecular chaperones that structurally remodel their protein substrates by threading them through the central channel of the ring-shaped oligomer (Puchades et al., 2020). Surprisingly, though, torsinA lacks pore loop consensus motifs, which are used by AAA+ proteins to engage and thread unfolded protein substrates through their central channel (Demircioglu et al., 2019). Furthermore, the substrate-binding domain normally found at the N-terminus of AAA+ proteins is replaced by a hydrophobic N-terminal domain (NTD) in torsinA that promotes its monotopic association with membranes and retention within the secretory pathway (Vander Heyden et al., 2011; Liu et al., 2003). Thus, while torsinA has been suggested to structurally remodel LINC complex proteins (Gerace, 2004; Saunders and Luxton, 2016), the protein substrates remodeled by torsinA within the NE have yet to be identified.

A significant advance in our mechanistic understanding of torsinA function within the NE was the identification of the INM protein lamina-associated polypeptide 1 (LAP1) and the ER/ONM protein luminal domain-like LAP1 (LULL1) as major torsinA-binding partners (Goodchild and Dauer, 2005). LAP1 and LULL1 are thought to activate torsinA, as the direct interaction of their luminal domains with torsinA stimulates its ability to hydrolyze ATP above negligible background levels *in vitro* (Zhao et al., 2013). The importance of LAP1 and LULL1 is further highlighted by the fact that the DYT1 dystonia-causing mutation (ΔE302/303, or ΔE) impairs the ability of torsinA to interact with or be stimulated by either protein (Naismith et al., 2009; Zhao et al., 2013). Structural studies revealed that the luminal domains of LAP1 and LULL1 possess an atypical AAA+ fold in which conserved arginine residues are positioned similar to the arginine finger that coordinates nucleotide hydrolysis in canonical AAA+ proteins (Demircioglu et al., 2019). Since torsinA lacks identifiable arginine fingers (Zhu et al., 2008), the torsinA-LAP1/LULL1 interaction was postulated to result in the formation of an alternating (torsinA-LAP1/LULL1)_3_ hetero-hexamer through which the arginine finger of LAP1 or LULL1 could coordinate ATP-hydrolysis with an adjacent torsinA (Brown et al., 2014; Sosa et al., 2014). However, the physiological relevance of this leading model of torsinA function is unknown, as the existence of the alternating (torsinA-LAP1/LULL1)_3_ hetero-hexamers in cells has yet to be demonstrated.

Two alternative models of torsinA function within the NE have been proposed. The first model postulates that torsins operate as a homo-hexamers that are disassembled when LAP1 or LULL1 stimulates their ability to hydrolyze ATP (Chase et al., 2017b; Goodchild et al., 2015). The second model hypothesizes that torsins function as helical filaments based on recent surprising cryo-electron microscopy results that show that torsinA forms helical filaments with a periodicity of 8.5 subunits per turn and a ∼4 nm diameter inner channel (Demircioglu et al., 2019). The ability of homo-oligomerization competent torsinA to induce the tubulation of membranes *in vitro* (Demircioglu et al., 2019) suggests that torsins promote interphase NPC biogenesis by mediating the asymmetric inside-out extrusion of the INM, which is required for this process in yeast and mammalian cells (Otsuka et al., 2016). Like the alternating (torsinA-LAP1/LULL1)_3_ hetero-hexamer model of torsinA function, neither homo-hexamers nor helical filaments of torsinA have been shown to exist within the NEs of living cells. Thus, to begin to understand the mechanism(s) underlying torsinA-dependent LINC complex and NPC assembly, the *in vivo* functional assembly states of this mysterious AAA+ protein within the NE need to be identified.

To address this goal, we used a powerful combination of fluorescence fluctuation spectroscopy (FFS), quantitative photobleaching analyses, and cell biological assays for torsin function. The knowledge gained by determining if torsinA regulates LINC complex and NPC assembly through a shared mechanism or divergent ones is essential for better understand the functional relationship between these important NE-spanning molecular bridges. This information is also indispensable for future efforts to rationally design novel therapeutic strategies and potential cures for DYT1 dystonia and other diseases caused by mutations in torsinA proteins, such as severe arthrogryposis (Kariminejad et al., 2017; Reichert et al., 2017; Iqbal et al., 2019; Siokas et al., 2017; Park et al., 2019).

## RESULTS

### TorsinA oligomerizes within the NE of living cells independently of LAP1 or LULL1

We used FFS to begin to distinguish between the three models of torsinA function within the NE as described above. FFS enables the characterization of fluctuating fluorescence signals generated by fluorescently labeled proteins passing through the optical observation volume, or point-spread function (PSF), of a two-photon microscope (Slaughter and Li, 2010). We recently developed FFS to be able to quantify the oligomeric state of NE proteins in living cells (Hur and Mueller, 2015a; Hennen et al., 2017; Hennen et al., 2018c; Hennen et al., 2018a; Hennen et al., 2018b; Hennen et al., 2020a). FFS measures the normalized brightness *b* of EGFP-tagged proteins (Chen et al., 2003), which indicates their average oligomeric state. For example, a monomeric and a dimeric EGFP-tagged protein have a *b* of 1 and 2, respectively. By measuring *b* vs. the number concentration *N*, which is the average number of EGFP-tagged proteins present within the two-photon PSF, a brightness titration is generated (Chen et al., 2003). Plotting *b* vs. *N* data reveals the concentration-dependent assembly state of a fluorescently labeled protein, which with increasing concentration approaches its limiting stoichiometry.

First, we performed FFS measurements in the NE of human osteosarcoma U2OS cells expressing a previously described and functional torsinA construct where EGFP was inserted immediately following the ER/NE lumen-targeting signal sequence (SS) of wild type (WT) torsinA (SS-EGFP-torsinA^WT^, (Goodchild and Dauer, 2004; Saunders et al., 2017)) (Fig. 1A). Plotting the *b* of SS-EGFP-torsinA^WT^ vs. *N* reveals a concentration-dependent increase in *b* with values between 2 and 3 for *N* > 100 (Fig. 1B). Since *b* vs. *N* represents a titration curve of protein binding in living cells, we expect that subtle differences of intracellular conditions between cells modulate the binding affinity, thereby leading to a range of titration curves. The scatter observed in the *b* values measured for SS-EGFP-torsinA^WT^ is consistent with the presence of cell-to-cell variability. However, we expect that this scatter would be reduced as *b* approaches the limiting stoichiometry of a protein complex, which has to be the same for all cells. Because such a reduction in scatter was not observed, the maximum *b* value shown in Fig. 1B may not be the true limiting stoichiometry of SS-EGFP-torsinA^WT^. Alternatively, the presence of endogenous non-fluorescent ‘dark’ torsinA could also reduce the measured *b* of SS-EGFP-torsinA^WT^. Since we did not see much of an impact of the presence of dark SUN1 or SUN2 on the measured *b* of SS-EGFP-tagged luminal domains of these proteins (Hennen et al., 2018c), we do not anticipate that the presence of dark torsinA will greatly influence our measurements of SS-EGFP-torsinA^WT^. Thus, we conclude that the limiting stoichiometry of SS-EGFP-tagged torsinA^WT^ must be at least that of a trimer.

**Figure 1.**
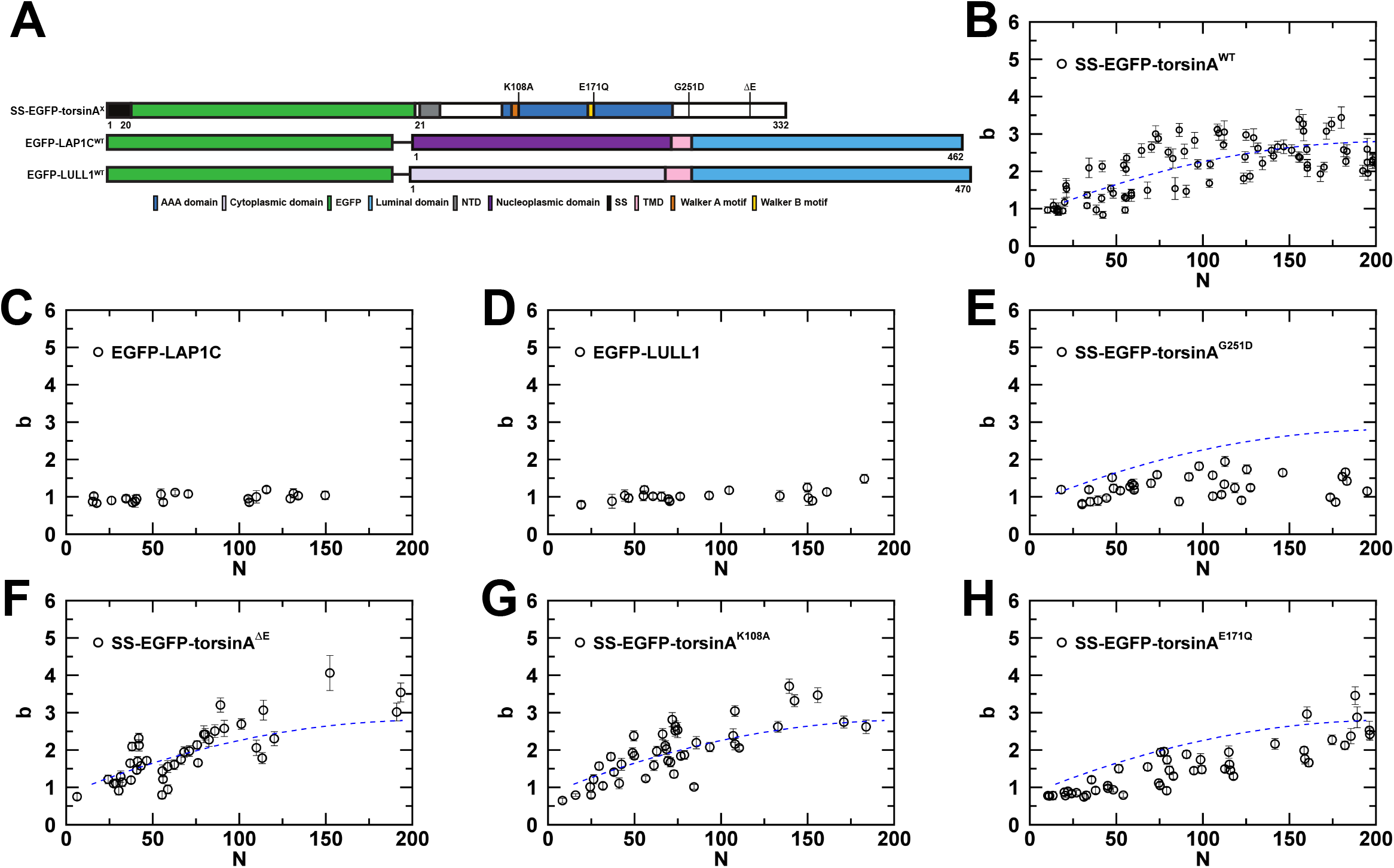
Analysis of torsinA, LAP1, and LULL1 oligomerization within the NE. A) Constructs used in this figure. B-H) Plots of *b* vs. *N* within the NE for the indicated constructs expressed in U2OS cells. The *b* vs. *N* trend of SS-EGFP-torsinA^WT^ is drawn as a blue dashed line (panel B) to guide the eye and provides a visual aid for comparison with the SS-EGFP-torsinA mutants shown in panels E-H.

The FFS results obtained for SS-EGFP-torsinA^WT^ within the NE of living cells with a brightness of ∼3 at high protein concentrations is consistent with the formation of an alternating hetero-hexamer of EGFP-tagged torsinA^WT^ with endogenous, dark LAP1 or LULL1. If this model were correct, we would expect to observe a brightness increase of EGFP-tagged LAP1 or LULL1 with concentration indicating the presence of hetero-complex formation with endogenous, dark torsinA^WT^. We therefore performed FFS measurements in the NEs of U2OS cells expressing a previously described and functional EGFP-tagged WT LAP1 construct that encodes the C isoform (EGFP-LAP1C^WT^) (Saunders et al., 2017) or an EGFP-tagged WT LULL1 construct (EGFP-LULL1^WT^) (Fig. 1C-D). Unlike SS-EGFP-torsinA^WT^, the *b* of either EGFP-LAP1C^WT^ or EGFP-LULL1^WT^ remained close to one and did not increase over the entire range of *N*, with *N* ≤ 200.

To further test the proposed alternating (torsinA-LAP1/LULL1)_3_ hetero-hexamer model of torsinA function within the NE, we next asked if the *N*-dependent increase in *b* observed for SS-EGFP-torsinA^WT^ was due to homotypic torsinA-torsinA or heterotypic torsinA-LAP1/LULL1 interactions. To do this, we used FFS to measure the limiting oligomeric states of SS-EGFP-torsinA encoding a mutation known to prevent the homo-oligomerization of torsinA *in vitro* (G251D) (Chase et al., 2017b) or the DYT1 dystonia-causing ΔE mutation that impairs the ability of torsinA to interact with or be activated by LAP1/LULL1 (Naismith et al., 2009; Zhao et al., 2013) (Fig. 1A). The *b* of SS-EGFP-torsinA^G251D^ was significantly reduced relative to the *b* measured for SS-EGFP-torsinA^WT^ across the entire range of *N*, with *N* ≤ 200 (Fig. 1E), supporting the homotypic model. In contrast, the *b* of SS-EGFP-torsinA^ΔE^ exhibited a *N-*dependent increase with values ranging between 2 to 3 for *N* > 100 that was indistinguishable from the *b* of SS-EGFP-torsinA^WT^ (Fig. 1F), suggesting that the disease-causing mutation does not disrupt the oligomerization of torsinA within the NE of living cells. These observations strongly argue that the oligomerization we observed with SS-EGFP-torsinA^WT^ is dominated by homotypic torsinA oligomerization rather than the formation of alternating (torsinA-LAP1/LULL1)_3_ hetero-hexamers in cells.

### Molecular requirements for torsinA oligomerization within the NE in living cells

Having established that homotypic torsinA-torsinA interactions are required for the *in vivo* oligomerization of torsinA within the NE, we hypothesized that torsinA may function as either a ring-shaped homo-hexamer or a higher-order homo-oligomeric filament. However, the maximum *b* value of SS-EGFP-torsinA^WT^ within the measured concentration range was ∼3 (Fig. 1B), which is smaller than expected for either a ring-shaped homo-hexamer or a higher-order homo-oligomeric filament of torsinA. Given that oligomerization and nucleotide-binding are often coupled in AAA+ proteins (Puchades et al., 2020) and *b* reflects the average oligomeric state of an EGFP-tagged protein (Chen et al., 2003), we hypothesized that the *b* values measured for SS-EGFP-torsinA^WT^ at *N* > 100 reflects the average nucleotide state of this construct in a particular cell. To test this hypothesis, we performed FFS measurements in the NEs of U2OS cells expressing previously described SS-EGFP-torsinA constructs harboring mutations known to inhibit ATP-hydrolysis (E171Q) (Goodchild and Dauer, 2004; Zhao et al., 2013) or predicted to prevent ATP-binding (K108A) (Goodchild and Dauer, 2004; Hanson and Whiteheart, 2005) (Fig. 1A). The brightness of both constructs increased with concentration with *b* values from 2 to 3 for *N* > 100 in a manner similar to what was observed for SS-EGFP-torsinA^WT^ (Figs. 1G-H). In contrast, the ability of SS-EGFP-torsinA^E171Q^ to oligomerize was reduced to the WT construct across the same concentration range.

We hypothesized that our inability to detect *b* values ≥ 6 for SS-EGFP-torsinA^WT^ within the NE may be related to how torsinA was tagged with EGFP in this construct. Goodchild and Dauer (2005) generated SS-EGFP-torsinA^WT^ by inserting EGFP immediately following the SS of torsinA (Fig. 2A) (Goodchild and Dauer, 2004), which is cleaved off the mature protein within the shared lumen of the ER/NE by the signal peptidase complex (Callan et al., 2007). Amino acids 26 through 43 of torsinA make the hydrophobic N-terminal domain (NTD), which is responsible for torsinA being a luminal monotopic membrane protein (Vander Heyden et al., 2011). Therefore, it is possible that the presence of EGFP at the N-terminus of torsinA has a negative impact on the ability of SS-EGFP-torsinA^WT^ to properly associate with nuclear membranes. Alternatively, torsinA was recently shown to undergo another proteolytic cleavage event that selectively removes amino acids 21 through 49 in response to ER stress and B cell stimulation (Zhao et al., 2016). This cleavage event results in the liberation of torsinA^49-332^ from the NTD-containing hydrophobic N-terminus. Thus, we hypothesized that our observation that the average oligomeric state of SS-EGFP-torsinA^WT^ at *N* > 100 was between a dimer and trimer may arise from the cleavage-dependent generation of two products: SS-EGFP-torsinA^21-49^ and torsinA^50-332^.

**Figure 2.**
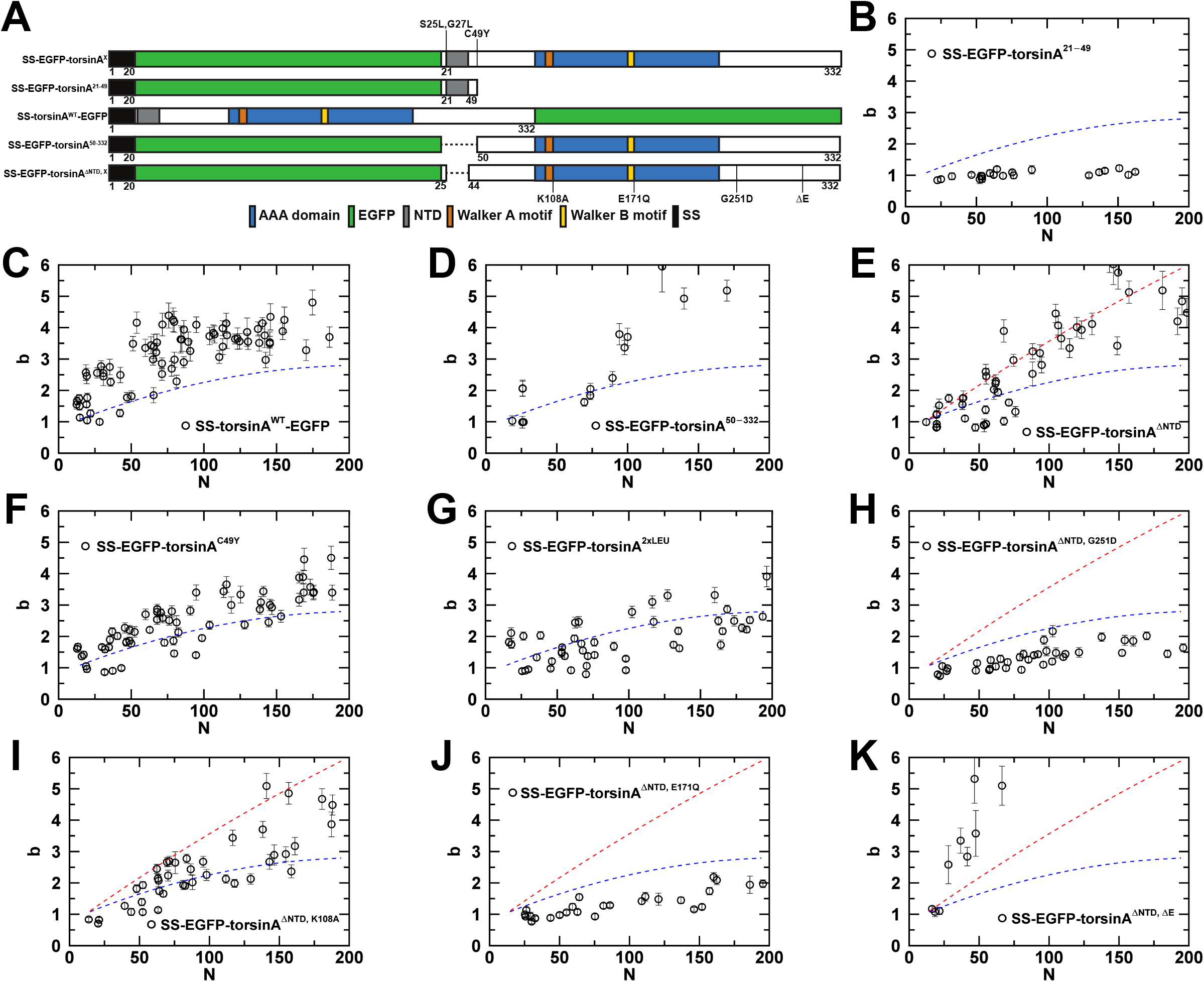
The oligomerization of torsinA within the NE is influenced by its association with the nuclear membranes. A) Constructs used in this figure. B-K) Plots of *b* vs. *N* within the NE for the indicated constructs expressed in U2OS cells. The blue and red dashed lines are visual guides for comparison with the *b* vs. *N* plot of SS-EGFP-torsinA^WT^ from Fig. 1B and SS-EGFP-torsinA^ΔNTD^ from panel E, respectively.

When we performed FFS in the NE of U2OS cells expressing SS-EGFP-torsinA^21-49^, the *b* of this construct did not increase significantly above a value of 1 across the entire range of *N* (Figs. 2A-B). This result indicates that the ability of torsinA to oligomerize must reside within amino acids 50 to 332. In addition, this result suggests that cleavage of the N-terminus only occurs for a minor population of SS-EGFP-torsinA^WT^, as the cleaved population would generate fluorescent SS-EGFP-torsinA^21-49^ and non-fluorescent (i.e. unobservable) torsinA^50-332^. The presence of a mixture of cleaved monomeric SS-EGFP-torsinA^21-49^ and uncleaved SS-EGFP-torsinA^WT^ in our FFS measurements would result in an effective reduction of the measured average oligomeric state as compared to a sample only containing uncleaved SS-EGFP-torsinA^WT^.

To further test the hypothesis that the measured *in vivo* oligomerization of SS-EGFP-torsinA^WT^ within the NE may be influenced by the proteolytic cleavage of the hydrophobic N-terminus of torsinA, we next used FFS to measure the brightness of a previously described torsinA^WT^ construct that contains monomeric EGFP fused to its C-terminus (SS-torsinA^WT^-EGFP; Fig. 2A) (Vander Heyden et al., 2009). As we observed for SS-EGFP-torsinA^WT^, the *b* of SS-torsinA^WT^-EGFP increased with *N* (Fig. 2C). At *N* values > 100; however, the average *b* of SS-torsinA^WT^-EGFP was > 3, which is larger than the *b* values measured for SS-EGFP-torsinA^WT^ across the same concentration range. This result suggests that the presence of EGFP at the N-terminus may limit the oligomerization of torsinA due to steric effects. Alternatively, it is possible that this result indicates that the limiting oligomeric state of SS-EGFP-torsinA^WT^ may be reduced by the cleavage of the first 49 amino acids of torsinA.

To begin to differentiate between these possibilities, we next used FFS to measure the oligomerization of a SS-EGFP-tagged torsinA construct that mimics the C-terminal cleavage product described above (SS-EGFP-torsinA^50-332^) (Fig. 2A). Like SS-EGFP-torsinA^WT^ and SS-torsinA^WT^-EGFP, the *b* of SS-EGFP-torsinA^50-332^ exhibited an *N*-dependent increase and at *N* values >100 continued to increase to *b* values ∼8 (Figs. 2D and S1A). This result supports the hypothesis that the first 49 amino acids of torsinA limit its oligomerization.

To determine if the higher-order oligomerization of SS-EGFP-torsinA^50-332^ were due to the presence of the NTD and therefore the ability of torsinA to associate with the nuclear membranes, we performed FFS in the NE of U2OS cells that expressed a torsinA construct from which we specifically deleted the NTD (SS-EGFP-torsinA^ΔNTD^) (Fig. 2A). The *b* of SS-EGFP-torsinA^ΔNTD^ behaved in a manner that was indistinguishable from the *b* of SS-EGFP-torsinA^50-332^ (Fig. 2E). Moreover, the *b* of SS-EGFP-torsinA^ΔNTD^ at *N* > 200 continued to increase up to ∼13 (Fig. S1B). Thus, we predict that inhibiting the proteolytic cleavage of the NTD-containing N-terminus or converting the NTD into a transmembrane domain would inhibit the oligomerization of torsinA. To inhibit the proteolytic cleavage of the N-terminus, we mutated cysteine 49 to a tyrosine in SS-EGFP-torsinA (SS-EGFP-torsinA^C49Y^; Fig. 2A), as the C49Y mutation was previously shown to prevent the N-terminal cleavage of torsinA in response to elevated levels of ER stress (Zhao et al., 2016). To convert the NTD into a transmembrane domain, we generated a construct that contains the previously described “2xLeu” mutation (Vander Heyden et al., 2011), where serine 25 and glycine 27 of torsinA are mutated to lysine (SS-EGFP-torsinA^2xLeu^; Fig. 2A).

The *b* values measured for SS-EGFP-torsinA^C49Y^ were significantly larger than those measured for SS-EGFP-torsin^WT^ (Fig. 2F), but were similar to those measured for SS-torsinA^WT^-EGFP (Fig. 2C), suggesting that the absence of proteolytic cleavage of the N-terminus of torsinA results in the same brightness produced by a combined mix of cleaved and uncleaved SS-torsinA^WT^-EGFP. In contrast, the *b* values measured for SS-EGFP-torsinA^2xLeu^ were indistinguishable from those measured for SS-EGFP-torsinA^WT^ (Fig. 2G). This result indicates that the conversion of the NTD into a transmembrane domain negatively impacts torsinA oligomerization in the NE.

To gain mechanistic insights into the higher-order oligomerization of torsinA constructs lacking the NTD, we performed FFS measurements on U2OS cells expressing SS-EGFP-torsinA^ΔNTD^ constructs harboring the G251D, K108A, E171Q, or ΔE mutations (Figs. 2H-K & S1C-D). Consistent with their effects on the oligomerization of SS-EGFP-torsinA^WT^, the G251D and E171Q mutations negatively impacted the oligomerization of SS-EGFP-torsinA^ΔNTD^, while the K108A mutation did not significantly alter its oligomerization. In stark contrast, the presence of the ΔE mutation resulted in a dramatic increase in the *b* of SS-EGFP-torsinA^ΔNTD^ starting at *N* values > 25. This observation indicates that the disease-causing ΔE mutation promotes the hyper-oligomerization of torsinA in the absence of the NTD.

### SS-EGFP-tagged strings of torsinA (SOTA) constructs exhibit significant immobilization within the NE of living cells

Given the shared molecular requirements between torsinA constructs containing or lacking the hydrophobic NTD, we wondered why we were unable to detect EGFP-tagged torsinA^WT^ oligomers that were greater than a tetramer. One possible explanation for this result is that the assembly of higher-order oligomers of torsinA on the nuclear membranes is a tightly regulated and therefore a low frequency event. To attempt to observe these rare higher-order torsinA oligomers on the nuclear membranes, we sought to perform FFS measurements in the NEs of U2OS cells expressing a SS-EGFP-tagged construct designed to increase the local concentration of torsinA by using flexible 20 amino acid linkers to covalently link together 5 torsinA^WT^ subunits to the C-terminus of SS-EGFP-torsinA^WT^, resulting in the formation of SS-EGFP-SOTA6 (Fig. 3A). This construct was designed following a previously described strategy where six subunits of the bacterial AAA+ protein ClpX were covalently linked together to study its function (Martin et al., 2005).

**Figure 3.**
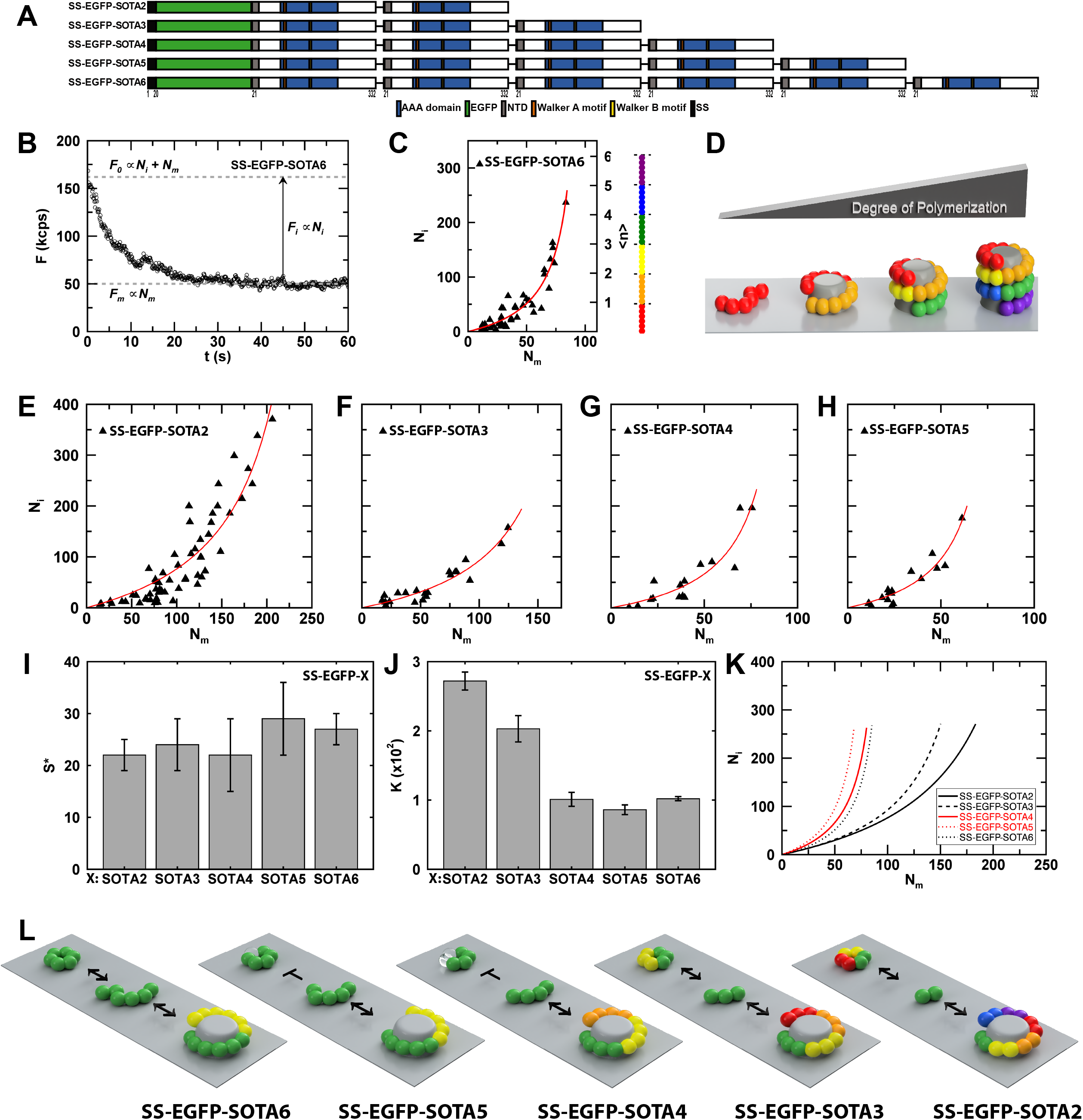
TorsinA immobilization and oligomerization at discrete sites within the NE. A) Constructs used in this figure. B) Representative fluorescent intensity trace (open circles) for SS-EGFP-SOTA6 measured within the NE showing a decrease from the peak value (*F*_0_) to a stable level (*F*_m_) with a decay amplitude of *F*_i_. The intensities *F*_m_ and *F*_i_ are proportional to the number of mobile (*N*_m_) and immobile (*N*_i_) molecules within the measured NE cross section. C) Plot of *N_i_* vs. *N_m_* for SS-EGFP-SOTA6 (triangles) measured within the NE together with a fit to the polymerization model (red solid line). The average degree of polymerization *n* is shown on the right axis and is graphically illustrated by the linear chain of SS-EGFP-SOTA6 molecules. Each SS-EGFP-SOTA6 molecule is drawn with a different color. D) Diagram showing SS-EGFP-SOTA6 oligomerization at a tubular seed site on the nuclear membrane (grey) as a function of the degree of polymerization. Individual SS-EGFP-SOTA6 constructs are represented by a different color. E-H) Plots of *N_i_* vs. *N_m_* for the indicated constructs measured within the NE with a fit to the polymerization model (red solid line). I) Density *S*\* of tubular seed sites per cross-sectional PSF area for all SS-EGFP-tagged SOTA constructs. J) Dissociation coefficient K for all SS-EGFP-tagged SOTA constructs. K) Fitted binding curves of the polymerization model for each of the SS-EGFP-SOTA constructs. L) Illustration of the formation of a helical SS-EGFP-SOTA polymer at a tubular seed site on the nuclear membrane (grey) for all SS-EGFP-SOTA constructs. The illustration also depicts the competing formation of a closed hexameric ring for all of the SS-EGFP-SOTA constructs except for SS-EGFP-SOTA4 and SS-EGFP-SOTA5. Individual SS-EGFP-SOTA constructs are represented by differently colored spheres. Cleared spheres represent missing torsinA molecules.

When we performed FFS measurements in the NE of cells expressing SS-EGFP-SOTA6, we observed the presence of a highly immobile subpopulation within this subcellular compartment. This is demonstrated by a significant reduction in fluorescence intensity of SS-EGFP-SOTA6 in the first few seconds of imaging (Fig. 3B). This observation is in stark contrast to the previously described SS-EGFP-torsinA^NTD-2xLeu^ construct (Hennen et al., 2017), which exhibited negligible photobleaching within the NE (Fig. S2A). The plots of the intensity of SS-EGFP-SOTA6 over time revealed an initial decay in its fluorescence amplitude, which eventually stabilized (Fig. 3B). Such intensity plots are indicative of the presence of a mixture of immobile and mobile fluorescently labeled proteins: the immobile population photobleaches while the mobile population diffuses quickly enough through the two-photon PSF to escape being photobleached (Skinner et al., 2008). Since FFS analysis in the presence of severe photobleaching introduces significant biases in the brightness that are difficult to correct for (Hur et al., 2014), the oligomeric state of SS-EGFP-SOTA6 could not be determined.

To gain insight into the assembly state of the immobile population of torsinA within the NE of living cells, we quantitatively analyzed the photobleaching of the SS-EGFP-SOTA6 construct within the NE of U2OS cells. The decay of the fluorescence amplitude depends on the initial intensity, which signals a concentration dependence of the mobile and immobile populations. Therefore, we quantified the number of mobile *N_m_* and immobile *N_i_* SS-EGFP-SOTA6 proteins in cells expressing varying amounts of this construct. The *N_m_* was determined from the mean fluorescent intensity measured following the completion of the photobleaching process, while the *N_i_* was determined by subtracting the final fluorescence intensity from the peak value, as explained in the Material and Methods (Fig. 3B). Plotting *N_i_* vs. *N_m_* for SS-EGFP-SOTA6 measured in the NE of U2OS cells revealed an unexpected convex curve (Fig. 3C). A typical binding titration curve relating the bound vs. free protein concentrations exhibits a concave curve with an initial linear increase that transitions into a plateau at high concentrations of free protein because of saturation of the binding reaction (Smith et al., 2015). In contrast, the concave upward curvature observed in the *N_i_* vs. *N_m_* plot for SS-EGFP-SOTA6 deviates from this expectation and cannot be explained by this simple binding equilibrium.

To account for the unusual curvature of the data, we had to develop a model based on SS-EGFP-SOTA6 polymerization at immobilization sites within the NE. This polymerization model is based on a mobile SS-EGFP-SOTA6 that diffuses along the luminal leaflets of the nuclear membranes until it comes into contact with a seed site. Once bound to the immobile seed site, polymerization growth occurs through the recruitment of additional mobile SOTA6 molecules with a constant binding affinity. The derivation is found in the Materials and Methods section, leading to Eq. 3, which relates *N_i_* and *N_m_*. The two free parameters of the model are the number of seed sites *S* within the cross-sectional area of the PSF and the dissociation coefficient *K*. This model of SS-EGFP-SOTA6 oligomerization is supported by the good fit of the *N_i_* vs. *N_m_* data for SS-EGFP-SOTA6 to Eq. 3 (red solid line, Fig. 3C) with *S* = 53 ± 6 and *K* = 102 ± 3. The degree of polymerization 〈*n*〉 describes the average number of SS-EGFP-SOTA6 molecules within the polymer, which is depicted in Fig. 3C on the right axis and was obtained by relating *N_i_* and *n* via Eq. 2.

The fluorescence data cannot provide information on the spatial conformation of the growing polymer at the NE membrane. However, a recent report by the Schwartz group observed that purified torsinA formed helical filaments that induced the tubulation of liposomes *in vitro* (Demircioglu et al., 2019). The interior of the torsinA filament is associated with the tubulated lipid membrane. Since the mean diameter of these membrane tubules was ∼20 nm, the authors estimated that the diameter of the membrane-bound filaments was ∼8.5 nm with ∼12.5 subunits per turn. Based on these results, we hypothesized that the immobilization seed site might also consist of about 12 torsinA-binding sites, which suggests a polymerization growth model as depicted in Fig. 3D, where two SS-EGFP-SOTA6 molecules define the initial seed side on the nuclear membrane. Further recruitment of SS-EGFP-SOTA6 units at the base of the cylindrical filament initiates vertical growth of the polymer complex that is accompanied by a simultaneous extrusion of the lipid membrane tube occupying the core of the protein filament. Since this model suggests that tubular membrane growth requires a base of about twelve torsinA proteins, it is sensible to define a new parameter *S*\* that characterizes the number of tubular seed sites per cross-sectional area of the PSF (0.17 ± 0.04 μm^2^). Since the fitted parameter *S* describes the number of SS-EGFP-SOTA6 seeds present within the PSF area and two such molecules defining a tubular seed site (Figs. S3A-B), we arrive at *S*\* = *S*/2 = 27 ± 3 sites per PSF area (Fig. 3I).

To further explore this helical polymerization hypothesis, we performed the same quantitative photobleaching experiments within the NE of U2OS cells expressing SS-EGFP-SOTA constructs containing, 2, 3, 4, or 5 covalently linked torsinA^WT^ subunits (SS-EGFP-SOTA2-5, Fig. 3A). Like SS-EGFP-SOTA6, the SS-EGFP-SOTA2-5 constructs were highly immobile within the NE (Figs. S2B-F) and their *N_i_* vs. *N_m_* curves have a concave-upward shape (Figs. 3E-H). Fits to the polymerization at immobile seed sites within the NE model described in Eq. 3 showed good agreement with each data set, recovering values of *S\** = 22 ± 3 and *K* = 272 ± 13 for SS-EGFP-SOTA2, *S\** = 24 ± 5 and *K* = 203 ± 19 for SS-EGFP-SOTA3, *S\** = 22 ± 7 and *K* = 101 ± 10 for SS-EGFP-SOTA4, and *S\** = 29 ± 7 and *K* =86 ± 7 for SS-EGFP-SOTA5 (Figs. 3I-J). The fitted S value was converted to *S*\* = *nS* / 12, where *n* represents the number of torsinA subunits for a given SS-EGFP-SOTA construct. Remarkably, the *S*\* values measured for each of the 5 SS-EGFP-SOTA constructs were identical within experimental uncertainty.

Plotting the fit curves obtained for each SS-EGFP-SOTA construct together revealed a trend in the data (Fig. 3K). Specifically, the fit curves for SS-EGFP-SOTA2, -SOTA3, and -SOTA6 follow a pattern where the larger number of torsinA subunits present within a SS-EGFP-SOTA construct results in an increasingly leftward shift of the fit curve towards lower *N_m_* values and represents an increased binding affinity. This observation indicates that the number of stabilizing interactions necessary for forming the helical filament increases with the number of torsinA subunits within the SS-EGFP-tagged SOTA construct. This explanation predicts that the fit curves of SS-EGFP-SOTA4 and SOTA-5 would fall between the fit curves of the SS-EGFP-SOTA3, and - SOTA6 constructs. However, this prediction is invalidated by the observation that the fit curves of SS-EGFP-SOTA4 and -SOTA5 are shifted to the left of the fit curve of SS-EGFP-SOTA6. Collectively, these results strongly suggest a model where torsinA may function within the NE via the assembly of a hexameric ring that cycles between open and closed states or as a higher-order helical oligomer. While additional subunits of torsinA could be added to an open hexameric ring resulting in the formation of a helical polymer, this would not be possible for a closed one. This model will be further addressed in considerable detail in the discussion.

### TorsinA immobilization is consistent with polymerization at discrete sites within the NE

To rule out the possibility that the insights into the mechanisms of torsinA oligomerization within the NE described above might represent SS-EGFP-SOTA construct-specific artifacts, we performed the same quantitative photobleaching analysis on the SS-EGFP-torsinA^WT^ construct expressed within the NE of U2OS cells. Despite being significantly reduced relative to the SS-EGFP-SOTA constructs, the fluorescent intensity vs. time plot of SS-EGFP-torsinA^WT^ exhibited non-negligible photobleaching (Fig. S2G). Plotting the *N_i_* vs. *N_m_* data for SS-EGFP-torsinA^WT^ revealed a curve similar to what was observed for the SS-EGFP-SOTA constructs (Fig. 4A). However, the amplitude of the curve achieved within the experimentally accessible concentration range was smaller than what was observed for any of the SS-EGFP-SOTA constructs, which made unconstrained fitting of the *N_i_* vs. *N_m_* data for SS-EGFP-torsinA^WT^ an ill-defined problem. Since we noticed that S*, the number of tubular seed sites per PSF area, was a conserved quantity (S* = 25 ± 3) among the 5 measured SS-EGFP-SOTA constructs (Fig. 3I), fitting of the *N_i_* vs. *N_m_* data was performed with S* fixed to 25, which recovered *K* = 1,090 ± 180 (red solid line, Fig. 4A). This result indicates that the same polymerization model used to describe the SS-EGFP-SOTA constructs was applicable to SS-EGFP-torsinA^WT^ (Fig. 4B), albeit with a dissociation constant of immobilization *K* that was significantly increased relative to the *K*’s of the SS-EGFP-SOTA constructs. The rightward shift of the fit curve towards higher *N_m_* values as compared to SS-EGFP-SOTA2 (black solid line, Fig. 4C) is consistent with our earlier observation that the number of torsinA subunits present within a SS-EGFP-tagged SOTA construct stabilizes the interactions responsible for forming the helical filament.

**Figure 4.**
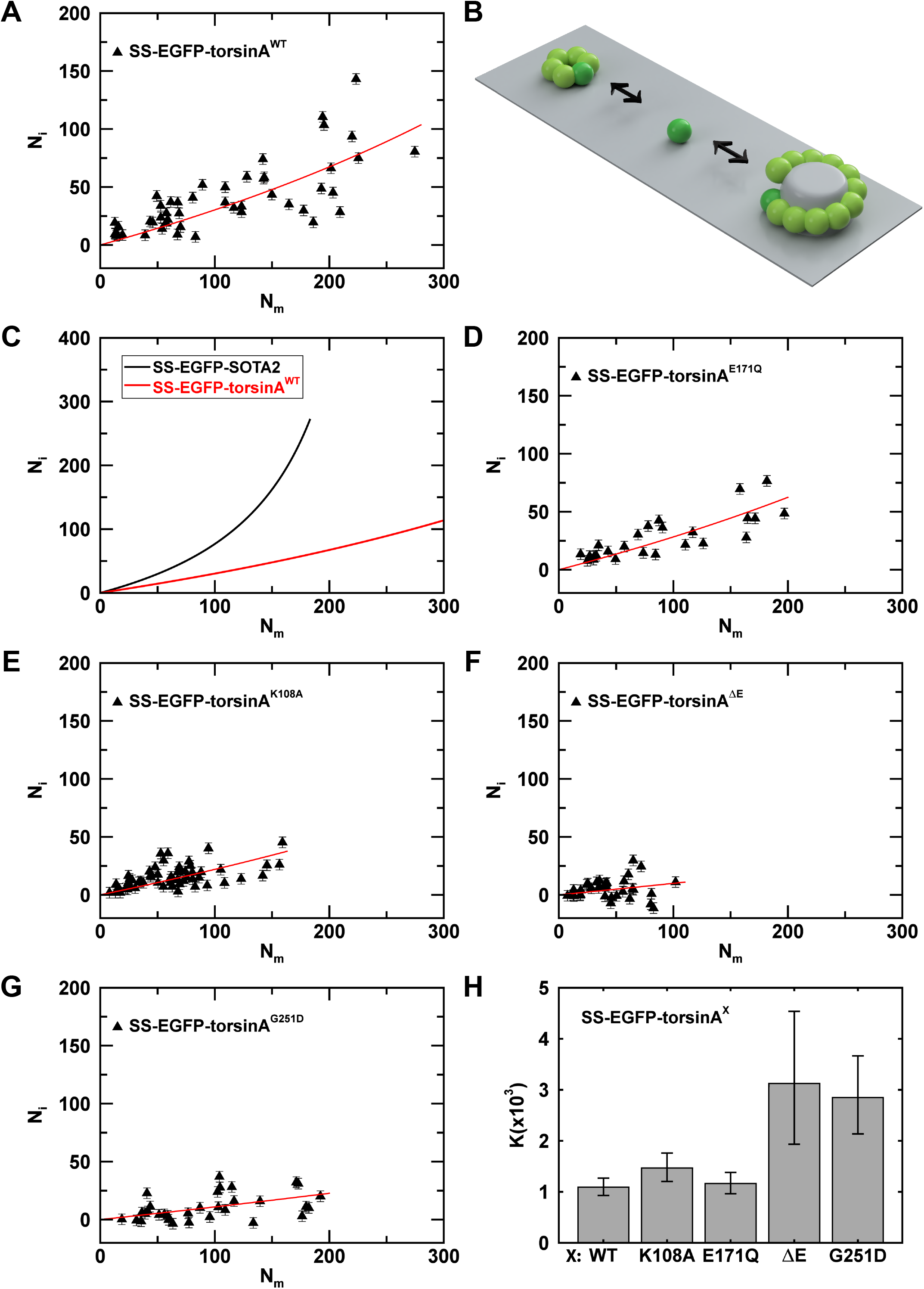
Immobilization of SS-EGFP-tagged WT and mutant torsinA constructs within the NE. A, D-G) Plots of *N_i_* vs *N_m_* for the indicated constructs measured within the NE of U2OS cells. B) Illustration of SS-EGFP-torsinA^WT^ forming a helical polymer at a tubular seed site with competing formation of a closed hexameric ring. Dark and light green spheres represent newly synthesized and older SS-EGFP-torsinA^WT^ molecules, respectively. C) Plot of the fitted binding curve for SS-EGFP-torsinA^WT^ and SS-EGFP-SOTA2. H) Bar plot of the average dissociation coefficient *K* for the indicated SS-EGFP-torsinA constructs together with the standard error of the mean.

Since the FFS measurements of the oligomerization of the SS-EGFP-tagged torsinA constructs expressed in the NEs of U2OS cells described above were performed on the mobile population of fluorescently labeled proteins (Hennen et al., 2017; Hennen et al., 2018c; Hennen et al., 2018a; Hennen et al., 2018b), we tested the effect of the G251D, K108A, E171Q, or ΔE mutations on the immobilization of SS-EGFP-torsinA by analyzing the photobleaching from the fluorescence intensity traces of these constructs. Small, but detectable levels of photobleaching were observed for all constructs. The *N_i_* vs. *N_m_* data obtained for each SS-EGFP-torsinA construct were fit by Eq. 3 with S* fixed to 25 (Figs. 4D-G), with the recovered dissociation coefficients shown in Fig. 4H together with SS-EGFP-torsinA^WT^ for comparison. While the K108A and E171Q mutations had no observable effect on the polymerization of SS-EGFP-torsinA, the ΔE and G251D mutation led to a larger mean value of K as compared to WT (Fig. 4H), suggesting a weakening of monomer binding to the filament. However, the experimental data (Fig. 4F) were insufficient to detect a statistically significant difference in the K between the ΔE mutation and SS-EGFP-torsinA (*p* = 0.1). In contrast, the K value of the G251D mutation differs significantly from WT (p = 0.02), indicating that the G251D mutation significantly weakens the formation of immobile torsinA filaments.

The decrease in fluorescence intensity due to photobleaching of the sample introduces biases in the brightness determination by FFS analysis (Hur and Mueller, 2015a). Unlike the pronounced intensity decrease of SS-EGFP-SOTA6 observed in the NE of U2OS cells (Fig. 3B), the amplitude of the intensity decay of SS-EGFP-torsinA^WT^ was small and only noticed upon close reexamination of the FFS data (Fig. S2A). Consequently, the bias in the reported brightness values for SS-EGFP-torsinA^WT^ (Fig. 1B) is expected to be minor. Additional tests were performed as described in the Material and Methods section to determine the magnitude of the bias for SS-EGFP-torsinA^WT^, its mutants, as well as EGFP-LAP1C^WT^ and EGFP-LULL1^WT^. These analyses found that brightness bias in these samples is entirely negligible (Fig. S4).

### Rearward nuclear positioning during centrosome orientation in fibroblasts polarizing for directional migration requires the hexamerization of torsinA and the cleavage of its hydrophobic N-terminus

Based on the results described above, we hypothesized that torsinA may function within the NE via the assembly of a hexameric ring that cycles between open and closed states or as a higher-order helical oligomer. To begin to test these hypotheses, we asked if the oligomerization-defective SS-EGFP-torsinA^G251D^ construct was able to rescue the defect in rearward nuclear positioning during centrosome orientation observed in *TorA*^-/-^ mouse embryonic fibroblasts (MEFs) polarizing for directional migration (Saunders et al., 2017). We previously demonstrated that expressing SS-EGFP-torsinA^WT^ in serum-starved wound-edge *TorA*^-/-^ MEFs that were stimulated with the serum component lysophosphatidic acid (LPA) restored their ability to orient their centrosomes and move their nuclei rearward to the levels observed in *TorA*^+/+^ MEFs. In contrast, the expression of SS-EGFP-torsinA^G251D^ was unable to rescue the ability of the *TorA*^-/-^ MEFs to orient their centrosomes or position their nuclei rearward (Figs. 5A-C).

**Figure 5.**
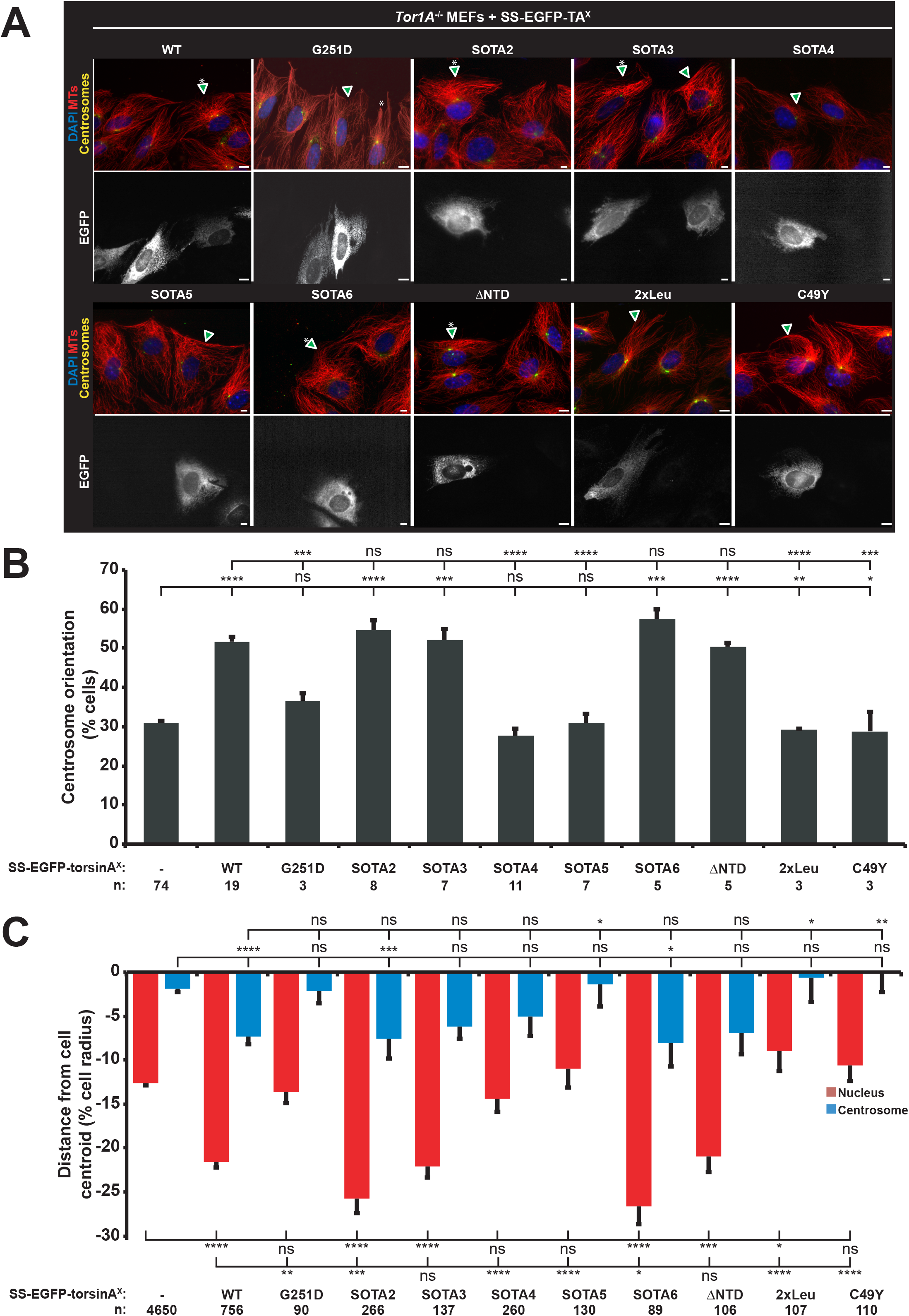
Rearward nuclear positioning during centrosome orientation requires the hexamerization of torsinA and the cleavage of its hydrophobic N-terminus. A) Representative epifluorescence images of centrosome orientation in *Tor1A*^-/-^ MEFs expressing the indicated cDNA constructs (arrowhead and bottom row of images). Asterisk: oriented centrosome. Scale bar: 5 μm. B) Average centrosome orientation in the cells described in A. C) Average positions of the centrosome and nucleus from the cells described in A. The cell center is defined as “0”. Positive and negative values represent positions that are toward the leading edge and cell rear, respectively. The error bars are standard error of the means. Two-tailed *t*-tests were used to calculate *p* values (\**p <* 0.05; \*\**p* < 0.01; \*\*\**p* < 0.001; \*\*\*\**p* < 0.0001; ns, not significant).

Next, we sought to determine the oligomeric assembly state required for torsinA-dependent centrosome orientation and rearward nuclear movement. Our above described quantitative photobleaching analyses predicted that the SS-EGFP-SOTA2, -SOTA3, and -SOTA6 constructs might be able to cycle between open and closed hexameric ring conformations, whereas neither SS-EGFP-SOTA4 nor -SOTA5 be able to assemble pseudohexamers. Like the SS-EGFP-torsinA^WT^ construct, the expression of SS-EGFP-SOTA2, -SOTA3, or - SOTA6 constructs was able to rescue centrosome orientation and rearward nuclear positioning in the *TorA*^-/-^ MEFs (Figs. 5A-C). Contrarily, neither the expression of SS-EGFP-SOTA4 nor -SOTA5 was able to rescue these processes in the *TorA*^-/-^ MEFs.

We then tested the hypothesis that the hydrophobic NTD of torsinA may regulate its function within this subcellular compartment by asking if expressing the SS-EGFP-torsinA^ΔNTD^ construct in the *Tor1A^-/-^* MEFs could rescue their impaired centrosome orientation or rearward nuclear positioning. Unexpectedly, we found that the expression of SS-EGFP-torsinA^ΔNTD^ was able to rescue these processes in the *Tor1A^-/-^* MEFs to levels that were statistically indistinguishable from the expression of SS-EGFP-torsinA^WT^ (Figs. 5A-C).

To explore the possibility that membrane-association negatively regulates these related torsinA-dependent functions, we asked if expressing SS-EGFP-torsinA^2xLeu^ in the *Tor1A^-/-^* MEFs could rescue their centrosome orientation and rearward nuclear positioning defects. Consistent with the idea that membrane-association inhibits torsinA-dependent rearward nuclear positioning during centrosome orientation in polarizing fibroblasts, we found that the expression of SS-EGFP-torsinA^2xLeu^ was unable to rescue these processes in the *Tor1A^-/-^* MEFs above the levels observed in control cells (Figs. 5A-C). We next asked if this cleavage event were required for torsinA-dependent rearward nuclear positioning during centrosome orientation by testing the ability of the expression of SS-EGFP-torsinA^C49Y^ to rescue these processes in the *Tor1A^-/-^* MEFs. We found that the levels of centrosome orientation and rearward nuclear positioning in the *Tor1A^-/-^* MEFs expressing the SS-EGFP-torsinA^C49Y^ construct were statistically indistinguishable from those observed in control cells (Figs. 5A-C).

### Interphase NPC biogenesis requires the Walker A motif of torsinA and higher-order oligomerization on nuclear membranes

Collectively, the results of our *Tor1A*^-/-^ MEF centrosome orientation and rearward nuclear positioning during centrosome orientation rescue experiments support a model where these related cellular processes require the ability of torsinA to assemble hexameric oligomers and the proteolytic removal of the NTD. To determine if this model could also explain how torsinA mediates another of its functions within the NE, we tested the ability of the same SS-EGFP-torsinA and -SOTA constructs used in the *Tor1A*^-/-^ MEF experiments to rescue the defect in interphase NPC biogenesis observed in the previously described *Tor1A/1B/2A/3A*-KO HeLa cell line (Laudermilch et al., 2016). Defective interphase NPC biogenesis can be visualized in the *Tor1A/1B/2A/3A*-KO HeLa cells by an accumulation of immunofluorescently labeled K48-linked Ubiquitin (K48-Ub) foci around the NE. This phenotype could be rescued by the expression of a torsinA^WT^ construct that contained a C-terminal hemagglutinin (HA) epitope tag (torsinA^WT^-HA) (Laudermilch et al., 2016). Consistent with this result, we found that expressing our SS-EGFP-torsinA^WT^ construct also rescued the interphase NPC biogenesis defect observed in the *TorA/B/2A/3A*-KO HeLa cells (Figs. 6A-C). Also consistent with the inability of torsinA-HA constructs harboring the G251D, ΔE, or ΔNTD mutations to rescue the interphase NPC biogenesis defect in these cells, we found that the expression our SS-EGFP-torsinA^G251D^, - torsinA^ΔE^, or -torsinA^ΔNTD^ constructs were not functional in in this assay (Figs. 6A-C). These results indicate that E302/303 and the NTD are critically important for torsinA-mediated interphase NPC biogenesis in HeLa cells. Despite being significantly reduced in its ability to rescue the interphase NPC biogenesis defect in HeLa relative to SS-EGFP-torsinA^WT^, the SS-EGFP-torsinA^G251D^ construct was clearly more functional than either the SS-EGFP-torsinA^ΔE^ or -torsinA^ΔNTD^ constructs.

**Figure 6.**
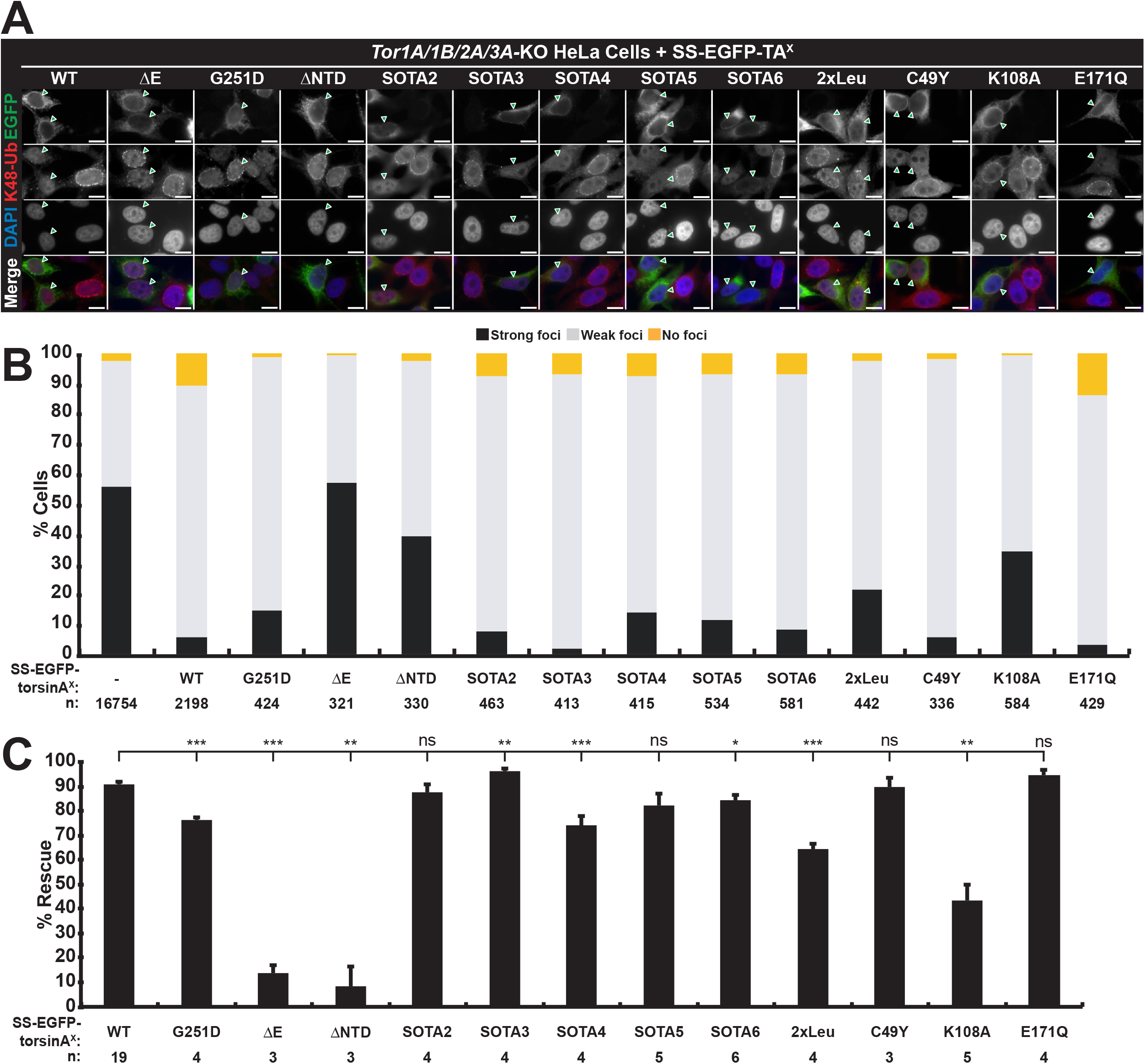
Interphase NPC biogenesis requires the Walker A motif of torsinA and higher-order oligomerization on nuclear membranes. A) Representative epifluorescence images of *Tor1A/1B/2A/3A*-KO HeLa cells expressing the indicated cDNA constructs (arrowheads) and stained with an anti-K48 ubiquitin antibody, an anti-EGFP antibody, and DAPI. Scale bar: 5 μm. B) Average percent of the cells described in A exhibiting K48 ubiquitin foci in the nuclear envelope. C) Average percent rescue of the K48 ubiquitin accumulation in the nuclear envelope in the cells described in A. The error bars are standard error of the means. Two-tailed *t*-tests were used to calculate *p* values (\**p <* 0.05; \*\**p* < 0.01; \*\*\**p* < 0.001; ns, not significant).

To further investigate the role of oligomerization during torsinA-dependent interphase NPC biogenesis, we tested the ability of our SS-EGFP-SOTA constructs to rescue this process in the *TorA/B/2A/3A*-KO HeLa cell line. While the SS-EGFP-SOTA4 construct was significantly reduced by ∼20% in its ability to rescue interphase NPC biogenesis in the *TorA/B/2A/3A*-KO HeLa cells relative to SS-EGFP-torsinA^WT^, the remaining 4 SS-EGFP-SOTA constructs were able to rescue this process to levels that were statistically indistinguishable from those observed in cells expressing SS-EGFP-torsinA^WT^ (Figs. 6A-C).

Given the requirement for the NTD during torsinA-mediated interphase NPC biogenesis in HeLa cells, we next asked if converting the NTD to a transmembrane domain or preventing the proteolytic cleavage of the NTD-containing N-terminus inhibited the ability of torsinA to function in this assay. We found that expressing the SS-EGFP-torsinA^2xLeu^ construct was significantly reduced by ∼30% in its ability to rescue the interphase NPC biogenesis defect in the *TorA/B/2A/3A*-KO HeLa cells relative to SS-EGFP-torsinA^WT^ construct, whilst the functionality of the SS-EGFP-torsinA^C49Y^ construct was statistically indistinguishable from this positive control (Figs. 6A-C).

Finally, we investigated the requirement for the ATP-binding Walker A and ATP-hydrolyzing Walker B motif during torsinA-dependent interphase NPC biogenesis by testing if the defects in this process observed in the *TorA/B/2A/3A*-KO HeLa cells could be rescued by expressing the SS-EGFP-torsinA^K108A^ or -torsinA^E171Q^, respectively. We previously demonstrated that neither of these constructs was capable of rescuing the rearward nuclear positioning or centrosome orientation defects observed in the *Tor1A*^-/-^ MEFs (Saunders et al., 2017). Consistent with these results, we found that the SS-EGFP-torsinA^K108A^ construct was significantly reduced by ∼50% relative to the SS-EGFP-torsinA^WT^ construct in its ability to promote interphase NPC biogenesis in the *TorA/B/2A/3A*-KO HeLa cells (Figs. 6A-C). In stark contrast with the results of the *Tor1A*^-/-^ MEF rescue experiments, the SS-EGFP-torsinA^E171Q^ construct was completely functional in this assay (Figs. 6A-C).

### The functions performed by torsinA are differentially sensitive to the fusion of EGFP to its C-terminus

As described above, the ability of the SS-EGFP-torsinA construct to rescue the centrosome orientation and rearward nuclear positioning defects in the *Tor1A*^-/-^ MEFs was inhibited by converting its NTD to a transmembrane domain with the 2xLeu mutation or preventing the proteolytic cleavage of its hydrophobic N-terminus with the C49Y mutation. While neither mutation impaired the ability of the SS-EGFP-torsinA construct to rescue the interphase NPC biogenesis defect in the *TorA/B/2A/3A*-KO HeLa cells, the deletion of the NTD did (Figs. 6A-B). Given these results, we hypothesized that rearward nuclear positioning during centrosome orientation required the dissociation of torsinA from the nuclear membranes, whereas interphase NPC biogenesis required the association of torsinA with the nuclear membranes. To further test this hypothesis, we asked if the SS-torsinA^WT^-EGFP construct could rescue centrosome orientation and rearward nuclear positioning in the *Tor1A*^-/-^ MEFs. This construct was selected based on our recent demonstration that the SS-torsinA^WT^-EGFP construct is more associated with the nuclear membranes than the SS-EGFP-torsinA^WT^ construct (Hennen et al., 2020b). We found that the SS-torsinA^WT^-EGFP construct was unable to rescue centrosome orientation or rearward nuclear positioning in the *Tor1A*^-/-^ MEFs (Figs. 7A-B). In contrast, the same construct was able to rescue interphase NPC biogenesis in the *TorA/B/2A/3A*-KO HeLa cells to levels that were statistically indistinguishable from SS-EGFP-torsinA^WT^ (Figs. 6C-D). These results provide further evidence to support a model whereby the functions performed by torsinA within the NE are sensitive to the fusion of EGFP to its C-terminus and therefore its ability to associate with the nuclear membranes.

**Figure 7.**
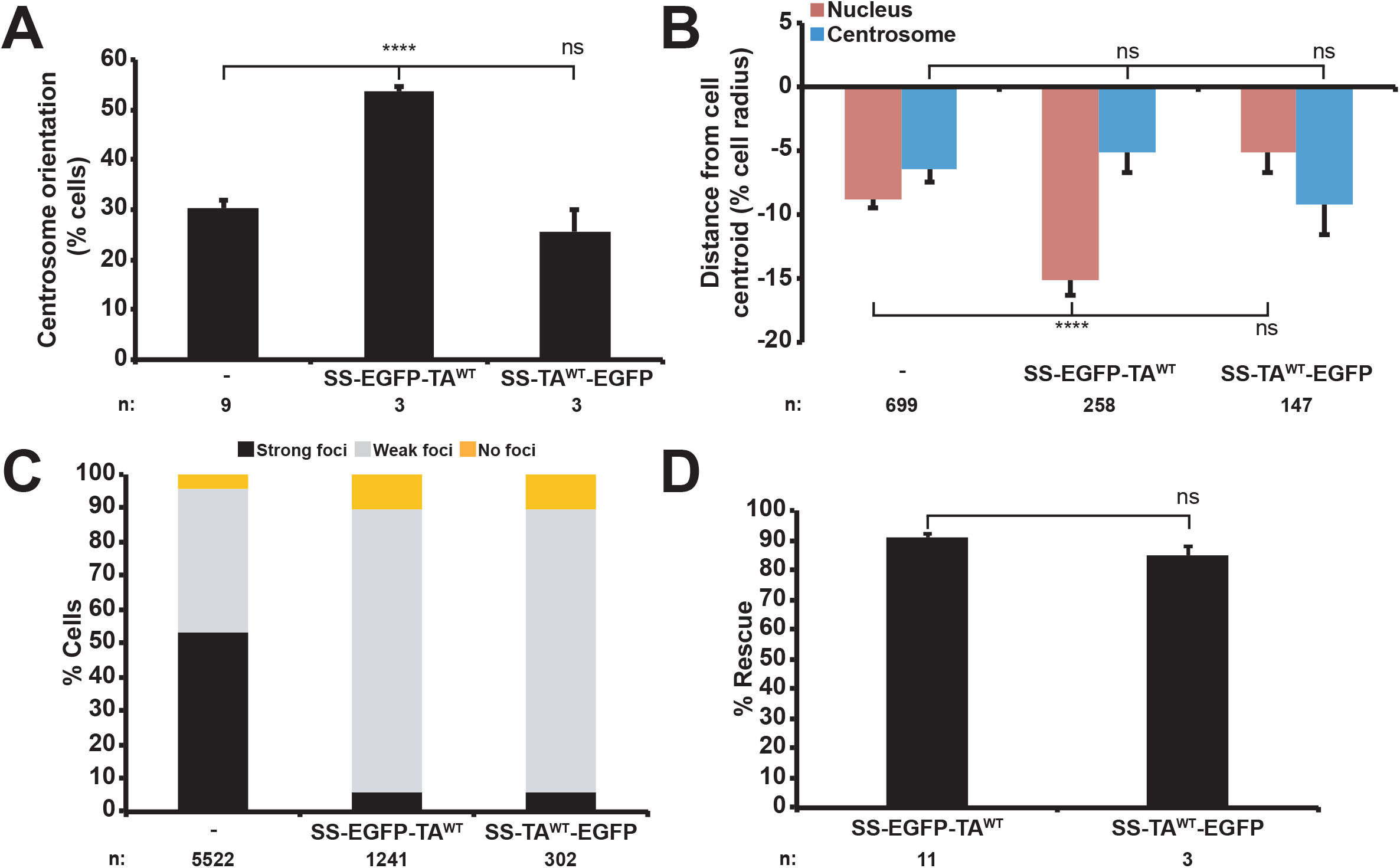
The functions performed by torsinA are differentially sensitive to the fusion of EGFP to its C-terminus. A) Average centrosome orientation in *Tor1A*^-/-^ MEFs expressing the indicated cDNA constructs. B) Average positions of the centrosome and nucleus from the cells described in A. C) Average percent of *Tor1A/1B/2A/3A*-KO HeLa cells expressing the indicated cDNA constructs that exhibit NE-localized K48 ubiquitin foci. D) Average percent rescue of the K48 ubiquitin accumulation in the nuclear envelope in the cells described in C. The error bars are standard error of the means. Two-tailed *t*-tests were used to calculate *p* values (\*\*\*\**p* < 0.0001).

## DISCUSSION

TorsinA is a luminal AAA+ protein that functions within the NE through poorly understood mechanisms. Here, we sought to begin to elucidate those mechanisms via an interdisciplinary approach that consisted of FFS, quantitative photobleaching analyses, and cell biological assays for torsinA function within the NE. Specifically, we wanted to investigate the physiological relevance of the three previously proposed models of torsinA function (i.e. the alternating (torsinA-LAP1/LULL1)_3_ hetero-hexamer, the homo-hexamer (torsinA)_6_, and the helical torsinA filament models). Our results identify the molecular requirements for torsinA oligomerization within the NE of living cells. In addition, our findings strongly suggest that interphase NPC biogenesis requires the ability of torsinA to associate with and remodel the INM by its ATP-dependent oligomerization into helical filaments that contain ≥ 12 subunits (Fig. 8A). Moreover, we propose that nuclear-cytoskeletal coupling requires the assembly of cleaved torsinA subunits lacking their membrane-associating NTD to assemble into homo-hexameric molecular chaperones that use the energy generated by ATP-hydrolysis to remodel the LINC complex (Fig. 8B). While our findings do not support the alternating (torsinA-LAP1/LULL1)_3_ hetero-hexamer model of torsinA function within the NE, we cannot exclude the possibility that these holoenzymes do exist within this subcellular compartment. Taken together, our results establish that torsinA acts via different assembly states to execute its distinct functions within the NE.

**Figure 8.**
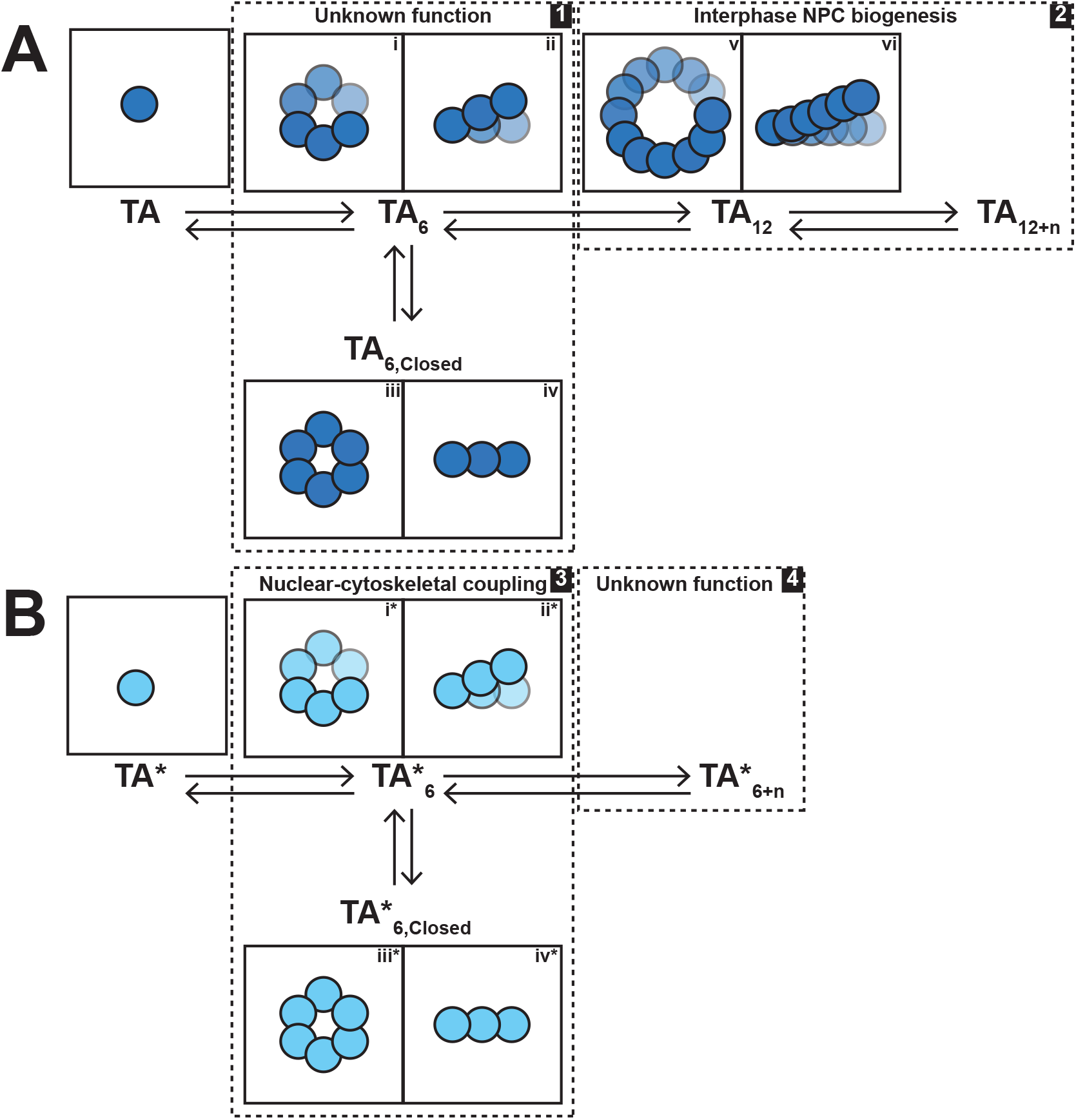
Model of differential torsinA function within the NE as a function of assembly state. A) Uncleaved torsinA (TA) molecules oligomerize into a hexamer that can be either a closed ring (TA_6, Closed_) or an open helical filament (TA_6_). We hypothesize that the transition between TA_6, Closed_ and TA_6_ may act as a power stroke for an as-of-yet unknown function performed on the nuclear membrane. The continued growth of TA_6_ into helical filaments containing ≥ 12 subunits (TA_12_ and TA_12+n_) at a tubular seed site on the INM enables the inside-out extrusion of the INM necessary for interphase NPC biogenesis through a mechanism that does not require ATP-hydrolysis. B) Proteolytically cleaved torsinA (TA*) molecules oligomerize into a hexamer that can be either a closed ring (TA*_6, Closed_) or an open helical filament (TA*_6_). We hypothesize that the transition between TA*_6, Closed_ and TA*_6_ may act as a power stroke for remodeling a protein complex required for LINC complex-dependent nuclear-cytoskeletal coupling during centrosome orientation in fibroblasts polarizing for migration. The continued growth of TA*_6_ into helical filaments that contain ≥ 6 subunits (TA*_6+n_) could perform an as-of-yet unknown function in the ER lumen and/or perinuclear space of the NE.

### The molecular requirements for the *in vivo* oligomerization of torsinA within the NE

The results of our FFS experiments performed within the NE of living cells demonstrate that torsinA oligomerizes within this subcellular compartment and identify the molecular requirements for this process. As expected from previously published *in vitro* biochemical and structural studies (Chase et al., 2017b; Demircioglu et al., 2019), we found that the ability of SS-EGFP-tagged torsinA to oligomerize within the NE was impaired by the presence of the E171Q or the G251D mutation. However, we were surprised to find that the K108A mutation did not have a major impact on the oligomerization of SS-EGFP-tagged torsinA in our experiments, as purified torsinA was unable to oligomerize in the absence of ATP (Chase et al., 2017b). Collectively, these results support the conclusion that the ATP-hydrolyzing Walker B motif is required for torsinA oligomerization, while the ATP-binding Walker A motif is dispensable. It should be noted that the contribution of the Walker A motif to the ability of torsinA remains to be experimentally determined. In addition, these results highlight the importance of homotypic torsinA-torsinA interactions for the *in vivo* oligomerization of torsinA within the NE.

Consistent with the hypothesis that neither LAP1 nor LULL1 appear to significantly contribute to the ability of torsinA to oligomerize within the NE, the *b* of either EGFP-LAP1C^WT^ or EGFP-LULL1^WT^ remained close to one and did not increase over the entire range of *N*, with *N* ≤ 200. This observation of monomeric EGFP-LAP1 or EGFP-LULL1 argues against the existence of an alternating (torsinA-LAP1/LULL1)_3_ hetero-hexamer *in vivo*, but definitive proof would require the use of dual-color FFS experiments performed in cells expressing fluorescently labeled torsinA and LAP1 or LULL1, which are technically challenging (Hennen et al., 2020a). Despite this lack of definitive proof, the results described above do not appear to support either the previously proposed alternating (torsinA-LAP1/LULL1)_3_ hetero-hexamer model of torsinA function within the NE.

Our FFS experiments also revealed that the presence of the hydrophobic NTD limits the ability of mobile SS-EGFP-tagged torsinA to oligomerize within the NE. We found that the *b* of SS-EGFP-torsinA^ΔNTD^ at *N* > 200 increase up to the highest *b* value that we measured, which was ∼13. We postulate that the molecular requirements of torsinA^ΔNTD^ are mostly shared with those used by torsinA^WT^, as the presence of the G251D or the E171Q mutations had similar negative effects on the oligomerization of either SS-EGFP-torsinA^WT^ or SS-EGFP-torsinA^ΔNTD^. In contrast, the presence of the K108A mutation slightly impaired the oligomerization of SS-EGFP-torsinA^ΔNTD^ at *N* > 100, while it had a negligible effect on the oligomerization of NTD-containing torsinA.

One potential limitation of our FFS experiments may be that they were performed in U2OS cells that express unlabeled endogenous torsinA, LAP1, and LULL1 proteins that may compete with our EGFP-tagged protein constructs, leading to a reduction in their brightness *b*. However, we previously demonstrated that the presence of unlabeled endogenous SUN proteins within the NE did not significantly impact the *b* of our labeled SUN protein constructs (Hennen et al., 2018c). The saturating value of *b* for these constructs remained unchanged and was approached after the exogenously expressed EGFP-tagged protein concentration exceeded the endogenous concentration as well as the KD value, identifying the stoichiometry of EGFP-tagged protein complexes (Chen and Müller, 2007). Since torsinA can interact with torsinB and torsin3A (Hewett et al., 2004; Naismith et al., 2009) and all torsin proteins can interact with either LAP1 or LULL1 (Jungwirth et al., 2010; Kim et al., 2010; Zhao et al., 2013), it is also possible that the *b* of our EGFP-tagged torsinA constructs may be influenced by the relative expression levels of these binding partners. It is important to note that the *b* values reported in this work represent a lower limit of the average stoichiometry of our EGFP-tagged protein constructs. Thus, future efforts where the oligomeric state of endogenous torsinA tagged with EGFP is measured by FFS in the absence or presence of known torsinA-binding partners are needed to better characterize the functional assembly states of torsinA within the NE.

### Working model of torsinA helical polymerization at discreet sites within the NE

The results of our quantitative SS-EGFP-tagged torsinA/SOTA construct photobleaching experiments strongly suggested that torsinA might be assembling into helical polymers at discreet sites within the NE. The simple polymerization model that we developed to attempt to explain the unusual curvature observed in the *N_i_* vs. *N_m_* plot for SS-EGFP-SOTA6 consists of a mobile SS-EGFP-SOTA6 molecule diffusing along the nuclear membranes of the NE that becomes immobilized upon its interaction with a seed site. After being bound to the seed site, polymerization growth can occur via the recruitment of additional mobile SS-EGFP-SOTA6 molecules with a constant binding affinity. The two free parameters of this model are the number of immobilization seed sites *S* present within the cross-sectional area of the PSF of our two-photon microscope and the dissociation coefficient *K*.

Given that the Schwartz laboratory had recently demonstrated that purified torsinA could form helical filaments that induced the tubulation of liposomes *in vitro* (Demircioglu et al., 2019), we hypothesized that the interior of the torsinA filament could be occupied by the presence of tubulated nuclear membrane. Taking into consideration the Schwartz laboratory’s estimation that the diameter of membrane-bound torsinA filaments was ∼8.5 nm with ∼12.5 subunits per turn (Demircioglu et al., 2019), we further hypothesized that the immobilization seed sites might also consist of ∼12 torsinA-binding sites. This would suggest that two SS-EGFP-SOTA6 molecules would define the initial seed site on the nuclear membrane and that the recruitment of additional SS-EGFP-SOTA6 molecules to the base of *S* would enable the vertical growth of the helical polymer while simultaneously extruding the nuclear membrane into the core of the polymer and consequently the perinuclear space. Given these assumptions, we proposed the definition of a new parameter *S\** that characterizes the number of immobilization seed sites for torsinA per cross-sectional area of the PSF of our two-photon microscope (0.17 ± 0.04 μm^2^). Thus, our data shows that *S\** of SS-EGFP-SOTA6 would be 27 ± 3 sites per PSF area.

Surprisingly, when we performed a similar quantitative analysis of our SS-EGFP-SOTA2, -SOTA3, - SOTA4, and -SOTA5 constructs and converted their fitted *S* vales into *S\** values using the equation *S*\* = *nS* / 12, where *n* is the number of torsinA subunits for a given SS-EGFP-SOTA construct, we found that all of the SS-EGFP-tagged SOTA constructs had *S\** values that were identical within experimental uncertainty. This result strongly suggests that the assembly of all the SS-EGFP-SOTA constructs occurs via the same assembly mechanism, resulting in an identical number of tubular seed sites. Based on the *in vitro* results from the Schwartz group (Demircioglu et al., 2019), we expect that each seed within the NE contains ∼12 torsinA-binding sites. Importantly, our quantitative photobleaching analyses of our SS-EGFP-tagged torsinA constructs support our conclusion that the above-described insights gained into the mechanisms of torsinA oligomerization are not SS-EGFP-SOTA construct-specific artifacts.

The deviation of the *N_i_* vs. *N_m_* curves of SS-EGFP-SOTA4 and -SOTA5 from the trend observed for SS-EGFP-SOTA2, -SOTA3, and -SOTA6 where the larger number of torsinA subunits present within a SS-EGFP-tagged SOTA construct causes an increasingly leftward shift of the fit curve towards lower *N_m_* values representing an increased binding affinity may be explained if another reaction were taking place in addition to helical growth that differentially affects SS-EGFP-SOTA4 and -SOTA5 from the other SS-EGFP-tagged SOTA constructs. Specifically, we hypothesize that torsinA forms both hexamers and helical filaments within the NE. This hypothesis is supported by recent structural studies, which show that a growing number of AAA+ proteins (e.g. ClpB, HSP104, katanin, Lon, and VPS4) operate as homo-hexamers or -dodecamers that cycle between open and closed ring conformations (Su et al., 2017; Gates et al., 2017; Zehr et al., 2017; Shin et al., 2020). Thus, SS-EGFP-SOTA6 would be expected to form helical filaments and homo-hexameric rings (Fig. 3L). Given that covalently linked homo-dimers or -trimers of ClpX can assemble into functional pseudohexamers by homo-trimerization and -dimerization, respectively (Martin et al., 2005), we predict that SS-EGFP-SOTA2 and - SOTA3 can also pseudohexamerize (Fig. 3L). Because SS-EGFP-SOTA6 is itself a single-chain pseudohexamer, we hypothesize that the SS-EGFP-SOTA2, -SOTA3, and -SOTA6 constructs may be able to cycle between open and closed hexameric ring conformations. In contrast, neither SS-EGFP-SOTA4 nor - SOTA5 would be predicted to be able to assemble pseudohexamers (Fig. 3L). Since the helical polymerization of torsinA can only occur on torsinA oligomers with an exposed subunit, a subpopulation of the SS-EGFP-SOTA2, -SOTA3, and -SOTA6 constructs would be in the closed ring conformation and unavailable for polymeric growth. This competing side reaction is absent for the SS-EGFP-SOTA4 and -SOTA5 constructs. Consequently, the polymeric growth of the SS-EGFP-SOTA2, -SOTA3, and -SOTA6 constructs would appear weaker than the polymeric growth of the SS-EGFP-SOTA4 and -SOTA5 constructs due to the added binding competition of the former set of constructs by forming closed, polymerization incompetent homo-hexameric rings. We currently do not know how the balance between an open and closed torsinA homo-hexamer is controlled in the perinuclear space. We hypothesize that a closed homo-hexamer, which is attainable on either a membrane or in solution, may provide a stabilizing interaction that preserves the ring structure. However, we speculate that a conformational change in torsinA must occur to draw membrane into the interior of the ring and that the ring must be sufficiently large enough to allow for membrane to fit into the center of the ring. Interestingly, the conformation of torsinA in the LULL1-bound state was shown to be incompatible with filament assembly due to steric hindrance (Demircioglu et al., 2019). Moreover, a rearrangement of the non-catalytic face of torsinA (residues 232-262) was observed, which was critical for the prevention of steric clashes with the nucleotide bound to the neighboring subunit. Perhaps this conformation change is related to the switch between open and closed homo-hexamers of torsinA. Future structural studies are clearly needed to be able to further understand the complex process of torsinA oligomerization within the NE.

In addition, a key question raised by our model of torsinA helical polymerization at discreet sites within the NE relates to the identity of the torsinA immobilization seed site. We hypothesize that these immobilization sites may consist of the INM protein Sad1/UNC-84 (SUN1) or the transmembrane nucleoporin POM121, which work together to promote the early steps of interphase NPC biogenesis (Talamas and Hetzer, 2011). In fact, SUN1 can interact with torsinA (Chalfant et al., 2019) and the depletion of SUN1 reduces the amount of NE-localized torsinA (Jungwirth et al., 2011). Moreover, POM121 is required for the formation of the NE blebs containing FG-nucleoporins and K48-linked ubiquitin found in cells lacking functional torsin ATPases (Rampello et al., 2020a). However, it should be noted that the density of the helical assemblies of torsinA in the NE is quite high at 160 ± 18 per μm^2^, while the density of NPCs in the NE of U2OS cells is ∼5.7 ± 2.7 per μm^2^ (Maimon et al., 2012). Alternatively, the torsinA immobilization seed sites might be defined by the presence of specific lipids and/or membrane curvature. We are currently exploring these different possibilities in our laboratory via a combination of cell biological and *in vitro* reconstitution experiments.

### TorsinA helical polymerization within the NE and interphase NPC biogenesis

TorsinA function is required for the interphase NPC biogenesis in *Caenorhabditis elegans*, humans, and mice (Laudermilch et al., 2016; Pappas et al., 2018; Rampello et al., 2020a; VanGompel et al., 2015). However, the mechanism underlying this proposed function remains unclear. Since purified torsinA can induce the tubulation of liposomes *in vitro* (Demircioglu et al., 2019), we hypothesized that torsinA helical polymerization may promote interphase NPC biogenesis by promoting asymmetric inside-out INM extrusion, which is critical for this process in yeast and mammalian cells (Otsuka et al., 2016). Consistent with previous reports (Chase et al., 2017b; Demircioglu et al., 2019; Laudermilch et al., 2016), we found that the presence of the G251D or the ΔE mutation as well as the deletion of the NTD impaired the ability of our SS-EGFP-tagged torsinA construct to rescue interphase NPC biogenesis in the previously described *Tor1A/1B/2A/3A*-KO HeLa cell line (Laudermilch et al., 2016). Thus, we conclude that G251-dependent homo-oligomerization, LAP1/LULL1-binding, and monotopic membrane-association are important for torsinA-mediated interphase NPC biogenesis, with LAP1/LULL1-binding and monotopic membrane-association being more important than G251-dependent homo-oligomerization.

We also demonstrated that each of our SS-EGFP-tagged SOTA constructs (i.e. SOTA2-6) was able to rescue interphase NPC biogenesis in the *Tor1A/1B/2A/3A*-KO HeLa cell line. While there is some difference in the significance of the ability of each of these constructs to rescue the interphase NPC biogenesis defect observed in the *Tor1A/1B/2A/3A*-KO HeLa cells, their performance was no worse than the performance of the SS-EGFP-torsinA^G251D^ construct in this assay. Based on these results and our above-described observation that each of our SS-EGFP-tagged SOTA constructs can assemble helical polymers within the NE via a similar mechanism.

Moreover, we provide evidence to support the hypothesis that the proteolytic cleavage of the NTD-containing N-terminus is dispensable for torsinA-mediate interphase NPC biogenesis. Specifically, we show that the NTD cleavage inhibiting mutation C49Y (Zhao et al., 2016) has a negligible effect on the ability of torsinA to promote interphase NPC biogenesis. Further support is provided by the ability of the SS-EGFP-tagged torsinA to rescue interphase NPC biogenesis in the *Tor1A/1B/2A/3A*-KO HeLa cells to ∼60% when its NTD is converted into a bona fide transmembrane domain via the 2xLeu mutation (Vander Heyden et al., 2011), as compared to the ∼90% rescue observed achieved by the SS-EGFP-torsinA^WT^ construct. This significant impairment in the rescue ability of the SS-EGFP-torsinA^2xLeu^ may arise from unanticipated steric effects induced by the 2xLeu mutation causing the NTD to become buried in the nuclear membrane. In the future, it would be interesting to determine if the 2xLeu mutation has a negative effect on the ATPase activity of purified torsinA reconstituted in liposomes.

Furthermore, the unimpaired ability of SS-EGFP-torsinA^E171Q^ to rescue interphase NPC biogenesis in the *Tor1A/1B/2A/3A*-KO HeLa cells strongly suggests that the ability of torsinA to hydrolyze ATP via its Walker B motif, and therefore the ability of torsinA to function as a canonical AAA+ molecular chaperone, is dispensable for interphase NPC biogenesis. On the other hand, the SS-EGFP-torsinA^K108A^ construct was significantly inhibited in its ability to rescue interphase NPC biogenesis in the *Tor1A/1B/2A/3A*-KO HeLa cells, suggesting that ATP-binding is critical for this process. Thus, we propose that ATP-binding and helical polymerization drive the extrusion of the INM necessary for interphase NPC biogenesis. Since the K108A mutation did not overtly impact the ability of torsinA to oligomerize within the NE as determined by our FFS or photobleaching experiments, ATP-binding may be important for inducing a conformational change in torsinA that is required for interphase NPC biogenesis. For example, a rearrangement of the non-catalytic face of torsinA is observed in the cryo-electron microscopy structure of helical torsinA^50-332^ that was proposed to avoid a steric clash with the bound nucleotide (Demircioglu et al., 2019).

### TorsinA homo-hexamerization and nuclear-cytoskeletal coupling during centrosome orientation

TorsinA function in the NE is also required for LINC complex-dependent nuclear-cytoskeletal coupling during centrosome orientation (Saunders et al., 2017). We previously showed that the defect in rearward nuclear movement during centrosome orientation observed in *Tor1A^-/-^* MEF polarizing for directional migration could be rescued by the expression of SS-EGFP-torsinA^WT^ but not with SS-EGFP-tagged torsinA constructs harboring the K108A, the E171Q, or the ΔE mutations. While the K108A and the ΔE mutations also impaired the ability of SS-EGFP-tagged torsinA to rescue the interphase NPC biogenesis defect seen in the *Tor1A/1B/2A/3A*-KO HeLa cells, this was not the case for the E171Q mutation. Thus, these results identified Walker B-mediated ATP-hydrolysis and LAP1/LULL1-binding as critically important for nuclear-cytoskeletal coupling during centrosome orientation.

Here, we demonstrate that G251-dependent oligomerization is required for nuclear-cytoskeletal coupling during centrosome orientation. While this result was somewhat similar to what we observed in the interphase NPC biogenesis assay, additional differences began to emerge regarding the ability of the SS-EGFP-tagged SOTA constructs to function in these two assays. Specifically, we found that the defective rearward nuclear positioning and centrosome orientation observed in the *Tor1A^-/-^* MEFs were rescued by SS-EGFP-SOTA2, -SOTA3, or -SOTA6, but not with SS-EGFP-SOTA4 or -SOTA5. We propose that these results could be potentially explained by the hypothesis that SS-EGFP-SOTA2, -SOTA3, and -SOTA6 may be able to switch between open and closed hexameric ring conformations, whereas neither SS-EGFP-SOTA4 nor - SOTA5 would be expected to pseudohexamerize. Thus, we hypothesize that nuclear-cytoskeletal coupling during centrosome orientation in polarizing fibroblasts requires the ability of torsinA to assemble hexameric oligomers. This may be related to the recently documented ability of AAA+ proteins like katanin to cycle between open spiral and closed ring conformations (Zehr et al., 2017). The cycling of the katanin hexamer was proposed to provide the power stroke for microtubule severing. Perhaps in this way, torsinA acts like a molecular disassemblase of nesprin-2G-containing protein complexes so to promote the proper mobility of nesprin-2G within the NE necessary for the assembly of transmembrane actin-associated nuclear lines, which are linear arrays of actin-binding LINC complexes that are required for rearward nuclear positioning during centrosome orientation in polarizing fibroblasts (Saunders et al., 2017). Since torsinA does not appear to have identifiable “pore loop” equivalents (Demircioglu et al., 2019), it is unlikely that torsinA hexamers will utilize a peptide threading mechanism involving a central pore. Therefore, to be able to better understand the molecular mechanism underlying torsinA-dependent nuclear-cytoskeletal coupling, future efforts should be dedicated to the *in vitro* reconstitution of torsinA-mediated LINC complex regulation.

In light of the findings described above, we hypothesize that torsinA-dependent nuclear-cytoskeletal coupling during centrosome orientation in fibroblasts polarizing for migration and interphase NPC biogenesis in HeLa cells operate via distinct molecular mechanisms. Further support for this hypothesis can be found in the differential effect that the deletion of the NTD as well as the 2xLeu and C49Y mutations have on the ability of torsinA to perform these two functions. For example, while the SS-EGFP-torsinA^ΔNTD^ cannot rescue interphase NPC biogenesis in *Tor1A/1B/2A/3A*-KO HeLa cells, it is able to rescue rearward nuclear positioning and centrosome orientation in *Tor1A^-/-^* MEFs to levels indistinguishable from SS-EGFP-torsinA^WT^. In addition, neither SS-EGFP-torsinA^2xLeu^ nor SS-EGFP-torsinA^C49Y^ was able to rescue rearward nuclear positioning and centrosome orientation in *Tor1A^-/-^* MEFs, while both constructs were at least partially functional in the interphase NPC biogenesis assay. These results strongly suggest that the NTD and therefore the ability of torsinA to associate with the nuclear membranes are dispensable for torsinA-dependent rearward nuclear positioning during centrosome orientation in fibroblasts. Moreover, they indicate that the proteolytic removal of the hydrophobic N-terminus of torsinA is critical for this process.

While we do not directly show that the hydrophobic N-terminus is proteolytically removed from torsinA within the NE, our FFS measurements of SS-torsinA^WT^-EGFP and SS-EGFP-torsinA^C49Y^ indirectly support this hypothesis because both form higher-order oligomers than SS-EGFP-torsinA^WT^. Intriguingly, the mitochondrial AAA+ protein Skd3 (i.e. human ClpB) was recently shown to exhibit enhanced disaggregase activity upon being cleaved by the mitochondrial inner membrane rhomboid protease presenilins-associated rhomboid-like (PARL) (Cupo and Shorter, 2020). The cleavage site for PARL in Skd3 is found within an autoinhibitory 35 amino acid hydrophobic stretch between the N-terminal mitochondrial targeting sequence and the ankyrin-repeat domain. Since Skd3 is a distant cousin of torsinA (Breakefield et al., 2001), we wonder if the presence of the NTD may autoinhibit the ability of torsinA to function as a disaggregase within the NE. Future investigations into this potential autoinhibitory mechanism will require the identification of the protease responsible for the proteolytic removal of the hydrophobic NTD of torsinA and the enigmatic protein substrate(s) remodeled by torsinA.

### Implications for DYT1 dystonia

It is important to consider how the results of this work may impact our mechanistic understanding and potential treatment of the diseases caused by the loss of torsinA function such as DYT1 dystonia and severe arthrogryposis (Ozelius et al., 1997; Kariminejad et al., 2017; Reichert et al., 2017). While the results of our FFS experiments revealed that the presence of the DYT1 dystonia-causing ΔE mutation had a negligible effect on the oligomerization of SS-EGFP-tagged torsinA within the NE, this was not the case for the SS-EGFP-torsinA^ΔNTD^ construct. Surprisingly, the ΔE mutation dramatically enhanced the oligomerization of SS-EGFP-torsinA^ΔNTD^ at *N* < 50, which is in agreement with previous *in cellulo* and *in vitro* reports that the ΔE mutation increases the amount of torsinA present in higher molecular mass species (Sosa et al., 2014; Zhao et al., 2013; Goodchild et al., 2015). Interestingly, the presence of the ΔE mutation was recently found to inhibit the ability of purified torsinA^50-332^ to assemble into helical polymers *in vitro* (Demircioglu et al., 2019). One potential explanation for these different effects regarding the impact of the ΔE mutation on torsinA oligomerization may be related to its ability to impair the interaction of torsinA with LAP1 or LULL1 and therefore be disassembled after hydrolyzing ATP (Chase et al., 2017b; Naismith et al., 2009; Zhao et al., 2013). Alternatively, the ΔE mutation may influence the oligomerization of torsinA by its ability to promote the formation of abnormal intermolecular disulfide bonds (Gonzalez-Alegre and Paulson, 2004; Torres et al., 2004). Thus, future studies will need to take into consideration the potential impact of LAP1/LULL1 and the redox environment of the ER and NE on torsinA oligomerization *in vivo*.

Since elevated levels of ER stress trigger the proteolytic removal of the hydrophobic NTD from the N-terminus of torsinA (Zhao et al., 2016) and ER stress is readily detected in the brains of DYT1 dystonia patients as well as fly and mouse models of DYT1 dystonia (Beauvais et al., 2016; Kim et al., 2015), we hypothesized that the hyper-oligomerization of SS-EGFP-torsinA^ΔNTD;ΔE^ may reveal potential insights into the mechanism underlying DYT1 dystonia pathogenesis. For instance, only 30-40% of ΔE mutation carriers develop DYT1 dystonia (Bressman et al., 2000). Perhaps individuals who have the ΔE mutation but do not develop DYT1 dystonia are those whose cells can effectively manage ER stress, while those who carry the ΔE mutation and suffer from DYT1 dystonia cannot. Because torsinA is a homeostatic regulator of the ER stress response (Chen et al., 2010; Hewett et al., 2003; Nery et al., 2011), we hypothesize that DYT1 dystonia pathogenesis in ΔE mutation carriers may require a second “hit” genetic mutation in other molecular modulators of ER stress.

While our studies were performed in human HeLa and U2OS cells as well as MEFs and DYT1 dystonia is a neurological movement disorder, the core molecular mechanisms underlying torsinA function within the NE are expected to be generally conserved across cell types. Thus, our findings may also be observed in neurons, which could have consequences for DYT1 dystonia pathogenesis. That being said, our demonstration that the E171Q and ΔNTD mutations can functionally uncouple torsinA-mediated nuclear-cytoskeletal coupling and interphase NPC biogenesis opens the door to exciting future experiments designed to test the differential contribution of defects in these two cellular functions to DYT1 dystonia pathogenesis using previously described fly, mouse, and worm models (Grillet et al., 2016; Liang et al., 2014; VanGompel et al., 2015). It will also be informative to test the effect of other DYT1 dystonia-associated mutations in *Tor1A* as well as the recently identified mutations in *Tor1A* associated with severe arthrogryposis (Kariminejad et al., 2017; Reichert et al., 2017; Iqbal et al., 2019; Siokas et al., 2017; Park et al., 2019). This information will be invaluable for the future rational design of novel treatment strategies for this debilitating disease.

### Conclusions and future directions

In summary, our data support the hypothesis that torsinA mediates nucleo-cytoskeletal coupling and interphase NPC biogenesis by adopting different functional assembly states within the NE. In addition, we demonstrate that the ability of torsinA to promote these two cellular processes is sensitive to the presence or absence of EGFP fused to its C-terminus. Moving forward, it will be exciting to test the effects of the E171Q and ΔNTD mutations on the following torsinA-dependent processes: ER-associated protein degredation (Esapa et al., 2007; Nery et al., 2011); nuclear egress of large ribonucleoprotein granules (Jokhi et al., 2013) and herpesvirus capsids (György et al., 2018; Maric et al., 2011; Maric et al., 2014; Turner et al., 2015; Hölper et al., 2020); nuclear membrane removal from mitotic chromatin (Luithle et al., 2020); and protein secretion (Hewett et al., 2007; Hewett et al., 2008; Torres et al., 2004). Finally, it will be important to begin to determine how torsinA oligomerizes within the peripheral ER, as torsinA is thought to be able to adopt multiple oligomeric states in this subcellular compartment (Chase et al., 2017a; Goodchild et al., 2015; Jungwirth et al., 2010). Since the ER is a highly dynamic organelle, we will explore the possibility of using a combination of imaging FFS and super-resolution microscopy (Bag and Wohland, 2014).

## Supporting information

Supplemental Figures Hur, Hennen et al., (2021) bioRxiv

## ACKNOWLEDGEMENTS

We thank members of the Luxton and Mueller laboratories for helpful discussions. Special acknowledgments go to S. D. Conner for the GM2163 *Escherichia coli strain* and the pcDNA3.1+ plasmid, W. T. Dauer for myc-LULL1^WT^, SS-EGFP-torsinA^WT^, SS-EGFP-torsinA^K108A^, SS-EGFP-torsinA^E171Q^, SS-EGFP-torsinA^ΔE302/303^, and the *Tor1a*^-/-^ MEFs; P.I. Hanson for SS-torsinA^WT^-EGFP; and Christian Schlieker for the control and TorA/B/2A/3A KO HeLa cells. This work was supported by the Dystonia Medical Research Foundation (G.W.G.L. and J.D.M.) and the NIH (AR007612 to C.A.S.; R01 GM129374-01 to GWGL; and R01 GM064589 to JDM).

## AUTHOR CONTRIBUTIONS

Conceptualization, G.W.G.L. and J.D.M.; Investigation, K.H.H., J.H., C.A.S., A.S., and C.M.M.; Construct generation, C.A.S., P.T.W., and G.W.G.L.; Formal Analysis, K.H.H., J.H., C.A.S., A.S., C.M.M., R.H.I., R.S.M., J.D.M., and G.W.G.L.; and Writing-Original Draft, K.H.H., J.H., G.W.G.L., and J.D.M..

## STAR METHODS

### CONTACT FOR REAGENT AND RESOURCE SHARING

Further information and requests for reagents and resources should be directed to and will be fulfilled by the Corresponding Authors, Joachim D. Mueller and G.W. Gant Luxton.

### EXPERIMENTAL MODEL AND SUBJECT DETAILS

FFS and quantitative photobleaching measurements were performed in low-passage number human U2OS bone osteosarcoma cells (authenticated by ATCC (Manassas, VA) and of unknown sex) that were cultured in DMEM with 10% bovine calf serum (Thermo Fischer Scientific, Waltham, MA). Wounded fibroblast monolayer experiments were performed in low-passage number *Tor1A*^-/-^ MEFs (previously described (Saunders et al., 2017) and of unknown sex). Monolayers of the *Tor1A*^-/-^ MEFs were cultured in DMEM with 15% bovine calf serum, serum starved for 3 days, wounded, and stimulated with 10 μM LPA (Avanti Polar Lipids, Inc., Alabaster, AL). Interphase NPC biogenesis assays were performed in *TorA/B/2A/3A*-KO HeLa cells (previously described (Laudermilch et al., 2016)) that were grown in DMEM containing 10% fetal bovine serum (Hyclone Laboratories, Logan, UT). All cell lines were grown at 37° C in 5% CO_2_.

### METHOD DETAILS

#### Reagents

DAPI was purchased from ThermoFisher Scientific. Restriction enzymes (REs) were either purchased from New England Biolabs (NEB, Ipswich, MA) or Promega (Madison, WI). Phusion DNA polymerase, T4 DNA ligase, and T4 PolyNucleotide Kinase (PNK) were also purchased from NEB. All other chemicals were from Sigma-Aldrich (St. Louis, MI) unless otherwise specified. GenJet *In Vitro* Transfection Reagent was purchased from SignaGen Laboratories (Rockville, MD). LPA was purchased from Avanti Polar Lipids, Inc. (Alabaster, AL). The QuikChange II site-directed mutagenesis kit was purchased from Aligent Technologies (Santa Clara, CA). The Wizard SV Gel and PCR Clean-Up System was from Promega. GeneJet Plasmid Midiprep Kit was from (ThermoFisher Scientific).

#### Cell culture and cDNA transfection

U2OS cells obtained from the American Type Culture Collection (ATCC, Manassas, VA) were cultured in Dulbecco’s Modified Eagle Media (DMEM) supplemented with 10% fetal bovine serum (Hyclone Laboratories, Logan, UT). The previously described *Tor1A*^-/-^ MEFs (Saunders et al., 2017) were cultured in DMEM with 15% bovine calf serum (ThermoFischer Scientific, Waltham, MA). We obtained the previously described *TorA*/*B*/*2A*/*3A* KO HeLa cells as a kind gift from Dr. Christian Schlieker (Yale University) (Laudermilch et al., 2016). These cells were cultured in DMEM with 10% fetal bovine serum. All cells were cultured using standard sterile technique and grown at 37 C° with 5% CO_2_. Transient transfections of cDNA constructs were performed using GenJet (SignaGen Laboratories, Rockville, MD) following the instructions provided by the manufacturer.

#### cDNA constructs

The generation of the EGFP-LAP1^WT^, SS-EGFP-torsinA^WT^, SS-EGFP-torsinA^K108A^, SS-EGFP-torsinA^E171Q^, SS-EGFP-torsinA^ΔE302/303^, and SS-torsinA^WT^-EGFP constructs was described previously (Goodchild and Dauer, 2004; Hennen et al., 2017; Hennen et al., 2018c; Naismith et al., 2004). The following constructs were generated during the course of this investigation using the primers described in Table 1. All constructs were confirmed by sequencing performed at the University of Minnesota Genomics Center. The standard *E. coli* DH5α strain was used for the majority of the molecular engineering described in this manuscript. However, the *E. coli* GM2163 strain, which is deficient in both *dam* and *dcm* genes, was used to propagate cDNA constructs that needed to be cut with Dam or Dcm-sensitive REs.

EGFP-LULL1^WT^ was generated by first PCR amplifying the cDNA encoding full-length LULL1 from the previously described myc-LULL1^WT^ construct (Goodchild et al., 2005) using the primers LULL1^WT^-F and LULL1^WT^-R. After purification, the resulting PCR product was digest alongside pEGFP-C1 with the REs *Sal*I and *Kpn*I, both of which were then gel purified and ligated together to create EGFP-LULL1^WT^. SS-EGFP-torsinA^G251D^, SS-EGFP-torsinA^ΔNTD^, SS-EGFP-torsinA^2xLeu^, and SS-EGFP-torsinA^C49Y^ were generated by PCR amplification using SS-EGFP-torsinA^WT^ as a template followed by PNK-ligase-*Dpn*I treatment of the purified resulting PCR products. SS-EGFP-torsinA^G251D^ was created using the primers torsinA^G251D^-F and torsinA^G251D^-R. SS-EGFP-torsinA^ΔNTD^ was generated using the primers torsinA^ΔNTD^-F and torsinA^ΔNTD^-R. SS-EGFP-torsinA^2xLeu^ was generated by two sequential PCR reactions. The first was performed using the primers torsinA^2xLeu^-F1 and torsinA^2xLeu^-R1, while the second was performed using torsinA^2xLeu^-F2 and torsinA^2xLeu^-R2 SS-EGFP-torsinA^C49Y^ was generated using the primers torsinA^C49Y^-F and torsinA^C49Y^-R.

SS-EGFP-torsinA^ΔNTD^ was created by QuikChange II site-directed mutagenesis using the primers torsinA^ΔNTD^-F and torsinA^ΔNTD^-R. We generated SS-EGFP-torsinA^ΔNTD, K108A^ by introducing the K108A mutation into SS-EGFP-torsinA^ΔNTD^ via PCR using the primers SS-EGFP-torsinA^ΔNTD,K108A^-F and SS-EGFP-torsinA^ΔNTD,K108A^-R followed by PNK-ligase-*Dpn*I treatment of the purified resulting PCR products. To generate SS-EGFP-torsinA^ΔNTD, E171Q^, SS-EGFP-torsinA^ΔNTD, G251D^, and SS-EGFP-torsinA^ΔNTD, ΔE^, we digested SS-EGFP-torsinA^ΔNTD^ and the corresponding mutated SS-EGFP-torsinA construct with *Bsr*GI. After gel purification, the resulting restriction enzyme digestion products were ligated together. SS-EGFP-torsinA^21-49^ and SS-EGFP-torsinA^50-332^ were both generated by PCR using SS-EGFP-torsinA^WT^ as the template and the primer pairs SS-EGFP-torsinA^21-49^-F and SS-EGFP-torsinA^21-49^-R and SS-EGFP-torsinA^50-332^-F and SS-EGFP-torsinA^50-332^-R, respectively. Following purification, the PCR products were separately subjected to PNK-ligase-*Dpn*I treatment.

To be able to create the SOTA constructs used in this manuscript, we first performed QuikChange II site-directed mutagenesis to mutate the *Nhe*I and *Spe*I RE cut sites found 5’ of the multiple cloning site (MCS) of pcDNA3.1+ following the manufacturer’s instructions. The *Nhe*I RE cut site was first mutated using the primers pSOTA*^Nhe^*^I^-F and pSOTA*^Nhe^*^I^-R, then the *Spe*I RE cut site was mutated using the primers pSOTA*^Spe^*^I^-F and pSOTA*^Spe^*^I^-R. The resulting product, pSOTA, was the vector in which the SOTA constructs were constructed. We then PCR amplified the cDNA encoding SS-EGFP-torsinA^WT^ minus the stop codon from SS-EGFP-torsinA^WT^ using the primers pSOTA.1/2/3-F and pSOTA.1-R. Following purification, the resulting PCR product was digested alongside pSOTA with the REs *AflI*II and *Apa*I. The digested PCR product and pSOTA were then gel purified and ligated together to create pSOTA.1.

Next, we performed a series of PCR reactions to insert the cDNA encoding a flexible linker sequence (L20*; ASGAGGSEGGGSEGGTSGAT) similar to the previously described L20 linker (ASGAGGSEGGGSEGGTS) that was used to build a similar construct that encodes covalently linked subunits of the bacterial AAA+ protein ClpX (Martin et al., 2005). The L20* cDNA was engineered such that it contained the cut sites for the REs *Bam*HI, *Kpn*I, *Sgr*AI, *Spe*I, *BspE1*, and *Apa*I, which were subsequently used to sub-clone the individual torsinA^WT^ subunits resulting in the assembly of the five SOTA constructs (SS-EGFP-SOTA_2_, SS-EGFP-SOTA_3_, SS-EGFP-SOTA_4_, SS-EGFP-SOTA_5_, and SS-EGFP-SOTA_6_) used in this work. Following each of the five engineered RE cut site was inserted six bases (TAGAAA) that served as both a stop codon and a spacer to enable proper RE-mediated DNA digestion. To do this, we first used the primers pSOTA.1/2/3-F and pSOTA.2-R to amplify the cDNA encoding SS-EGFP-torsinA. The resulting PCR product, which contains five silent mutations at the 3’ end of the cDNA encoding torsinA^WT^ (now referred to as torsinA^WT*^) in order to reduce the annealing temperature of the primers pSOTA.2-R and pSOTA.3*-R as well as half of the cDNA encoding L20*, was purified and digested alongside pSOTA.1 using the REs *AflI*II and *Apa*I. Following gel purification, the digested PCR product and pSOTA.1 were ligated together to create pSOTA.2. The cDNA encoding SS-EGFP-torsinA^WT*^ and half of L20* was PCR amplified from pSOTA.2 using the primers pSOTA.1/2/3-F and pSOTA.3*-R. The resulting PCR product, which encoded SS-EGFP-torsinA^WT*^-L20*, was digested alongside pSOTA.1 with the REs *AflI*II and *Apa*I. The digested PCR product and pSOTA.1 were then gel purified and ligated together to create pSOTA.3. To correct for a reading frame shift caused by the fact that the *Sgr*AI cut site encoded in the pSOTA.2-R primer that was incorrectly designed, PCR was performed using pSOTA.3 as a template and the primers pSOTA.1/2/3-F and pSOTA.3*-R. The purified PCR product and pSOTA.1 were then digested with the REs *AflI*II and *Apa*I. Afterwards, the digested PCR product and pSOTA.1 were gel purified and ligated together to create pSOTA.3*.

Having successfully generated pSOTA.3*, we next sought to create an entry vector (pSOTA_Entry_) into which we could sub-clone and sequence the PCR amplified torsinA subunits that would then be used to build the five SOTA constructs described here. To do this, we deleted SS-EGFP-torsinA^WT*^ from pSOTA.3* by performing QuikChange II site-directed mutagenesis using the primers pSOTA_Entry_-F and pSOTA_Entry_-R. This deletion leaves the appropriate parts of the L20* linker and the RE cut sites used for insertion into the construct. Next, we created pSOTA_Entry_ constructs containing the cDNA encoding torsinA^WT*^ flanked by DNA sequences encoding the L20* and RE cut sites necessary to sequentially assemble the SOTA constructs used in this manuscript. To create the pSOTA_Entry_torsinA_2_^WT*^, pSOTA_Entry_torsinA_3_^WT*^, pSOTA_Entry_torsinA_5_^WT*^, and pSOTA_Entry_torsinA_6_^WT*^ constructs we PCR amplified the cDNA encoding torsinA^WT*^ using the following primer pairs TorsinA_2_-F and TorsinA_2_-R; TorsinA_3_-F and TorsinA_3_-R; TorsinA_5_-F and TorsinA_5_-R; and TorsinA_6_-F and TorsinA_6_-R, respectively. To create the pSOTA_Entry_torsinA_4_^WT*^ construct, we performed nested PCR using the primer pairs TorsinA_4_.1-F and TorsinA_4_.1-R, and TorsinA_4_.2-F and TorsinA_4_.2-R. Following PCR amplification, the PCR products were purified and digested alongside pSOTA_Entry_ using the appropriate REs, as indicated in Table 1. The digested PCR products and pSOTA_Entry_ constructs were then gel purified and ligated together.

To assemble SS-EGFP-SOTA2, we digested pSOTA.3* and pSOTA_Entry_torsinA_2_^WT*^ with the REs *Bam*HI and *Kpn*I. Both digested pSOTA.3* and pSOTA_Entry_torsinA_2_^WT*^ were then gel purified and ligated together. To assemble SS-EGFP-SOTA3, SS-EGFP-SOTA2 and pSOTA_Entry_torsinA_3_^WT*^ were digested with the REs *Kpn*I and *Sgr*AI. The digested SS-EGFP-SOTA2 was then treated with CIP and gel purified alongside the digested pSOTA_Entry_torsinA_3_^WT*^, which were subsequently ligated together. To assemble SS-EGFP-SOTA4, SS-EGFP-SOTA3 and pSOTA_Entry_torsinA_4_^WT*^ were digested with the REs *Sgr*AI and *Spe*I. The digested SS-EGFP-SOTA3 was then treated with CIP and gel purified alongside the digested pSOTA_Entry_torsinA_4_^WT*^, which were subsequently ligated together. To assemble SS-EGFP-SOTA5, SS-EGFP-SOTA4 and pSOTA_Entry_torsinA_5_^WT*^ were digested with the REs *Spe*I and *Bsp*EI. The digested SS-EGFP-SOTA4 was then treated with CIP and gel purified alongside the digested pSOTA_Entry_torsinA_5_^WT*^, which were subsequently ligated together. To assemble SS-EGFP-SOTA6, SS-EGFP-SOTA5 and pSOTA_Entry_torsinA_6_^WT*^ were digested with the REs *Bsp*EI and *Apa*I. The digested SS-EGFP-SOTA5 was then treated with CIP and gel purified alongside the digested pSOTA_Entry_torsinA_6_^WT*^, which were subsequently ligated together.

#### Wounded fibroblast monolayer assay

*Tor1A*^-/-^ MEFs were grown on #1.5 coverslips until they reached ∼70% confluence, when they were starved of serum for three days. The serum-starved fibroblast monolayers were then wounded and the nuclei of wound-edge cells were injected with cDNA constructs that were purified using the Plasmid Midi Kit (Qiagen, Valencia, California) at concentrations between 5-30 μg/ml. Injections were performed on a Nikon (Melville, New York) TS100 microscope equipped with a CFI Achro LWD ADL 20X/NA0.4 WD 3.0mm objective and a Nikon Narishige NT-88 Manipulator set. After allowing the injected cDNA constructs to express for 2 hours at 37 C° with 5% CO_2_, the cells were stimulated with 10 µM LPA and returned to the incubator for an additional 2 hours before being fixed in −20° C methanol for 10 minutes.

The fixed cells were rehydrated in 1X Tris-buffered saline for 5 minutes then incubated for 30 minutes at room temperature in 10% normal goat serum (ThermoFisher Scientific) containing dilutions of the following primary antibodies: 1:200 anti-EGFP Living Colors A. v. mouse monoclonal antibody (JL-8, Clontech Laboratories, Inc., Mountain View, CA); 1:300 anti-pericentrin rabbit polyclonal antibody (PRB-432C, Covance, Princeton, NJ); and 1:50 anti-tyrosinated α-tubulin rat monoclonal antibody collected in house from supernatant generated by hybridomas (YL1/2, European Collection of Animal Cell Cultures, Salisbury, UK). The stained coverslips were washed at room temperature for 5 minutes thrice in 1x Tris-buffered saline, then incubated for 30 minutes at room temperature in 10% normal goat serum containing DAPI (Life Technologies, Calsbad, CA) and 1:200 dilutions of the following secondary antibodies: AlexaFluor488-conjugated goat anti-mouse polyclonal antibody; AlexaFluor555-conjugated goat anti-rat polyclonal antibody; and AlexaFluor650-conjugated goat anti-rabbit polyclonal antibody. The secondary antibodies were purchased from Jackson ImmunoResearch Laboratories (West Grove, PA) or Life Technologies. Following this last incubation, the stained coverslips were again washed at room temperature for 5 minutes thrice in 1x Tris-buffered saline. Finally, the stained coverslips were mounted on glass slides using Fluoromount (Thermo Fischer Scientific) and stored at 4 C° until they were ready to be imaged.

#### Quantification of centrosome orientation

To quantify centrosome orientation, a fibroblast is divided by drawing a “v” that begins and ends at the left- and right-most wound-edge, respectively. The middle of the “v” is defined by the center of the nucleus. The cell is scored as having an oriented centrosome if the centrosome resides within the “v” between the nucleus and the wound-edge. Since the “v” occupies ∼1/3 of the cell, ∼33% of wound-edge cells will have an oriented centrosome. Experiments were repeated ≥ 3 times.

#### Quantification of nucleus and centrosome position

To quantify nuclear positioning, fluorescence images of the stained cells described above were pseudocolored, combined and aligned such that the wound-edge was parallel to the x-axis using ImageJ. Custom-made MATLAB software (MathWorks, Natick, MA) was then used to calculate the cell centroid and equivalent radius of the wound-edge cells, which were outlined by hand. Segmentation was used to identify the centroid of the nucleus and therefore the position of the organelle. A vector representing the distance from the nuclear centroid to the cell centroid was drawn and resolved into x and y coordinates (parallel and perpendicular to the leading edge, respectively). Measurements were normalized to cell size to allow for comparison between cells. Only the y-coordinate was used in plots as the x-coordinate (position of the nuclear centroid or centrosome along the x-axis) did not change significantly. The difference between the cell centroid and the nucleus centroid or centrosome was then divided by the radius to determine the percentage of the cell radius the nucleus traveled.

#### K48-linked ubiquitin NE accumulation assay

*TorA*/*B*/*2A*/*3A* KO HeLa cells were grown on No. 1.5 glass coverslips (#152222, ThermoFischer Scientific) for a day before being transfected. The transfected cDNA constructs were then allowed to express for 18 hours and were subsequently fixed at room temperature for 20 minutes in 4% paraformaldehyde followed by 10 minutes of permeabilization in 0.1% tritonX-100 at room temperature as previously described (Chase et al., 2017b). Next, the fixed cells were rehydrated in 1X Tris-buffered saline for 5 minutes then incubated for 30 minutes at room temperature in 10% normal goat serum (ThermoFisher Scientific) containing dilutions of the following primary antibodies: 1:200 anti-EGFP Living Colors A. v. mouse monoclonal antibody (JL-8, Clontech Laboratories); and 1:500 anti-K48-linked ubiquitin rabbit monoclonal antibody (AB_11213655; Merck Millipore, Burlington, MA). The stained coverslips were washed at room temperature for 5 minutes thrice in 1x Tris-buffered saline, then incubated for 30 minutes at room temperature in 10% normal goat serum containing DAPI (Life Technologies, Calsbad, CA) and 1:200 dilutions of the following secondary antibodies: AlexaFluor488-conjugated goat anti-mouse polyclonal antibody; and AlexaFluor555-conjugated goat anti-rabbit polyclonal antibody. The secondary antibodies were purchased from Jackson ImmunoResearch Laboratories (West Grove, PA) or Life Technologies. Following this last incubation, the stained coverslips were again washed at room temperature for 5 minutes thrice in 1x Tris-buffered saline. Finally, the stained coverslips were mounted on glass slides using Fluoromount (Thermo Fischer Scientific) and stored at 4 C° until they were ready to be imaged.

#### Quantification of the K48-linked ubiquitin NE accumulation assay

Images of fixed and immunofluorescently stained *TorA*/*B*/*2A*/*3A* KO HeLa cells were assessed as strongly (≥ 20 foci), weakly (1-20 foci), or not (0 foci) exhibiting K48 foci in their nuclear envelopes, as previously described (Chase et al., 2017b). The percent rescue of the K48-linked ubiquitin accumulation phenotype was calculated using the following equation: Percent rescue = ((Percent of cells strongly exhibiting K48 foci in the controls - Percent of cells strongly exhibiting K48 foci in cells expressing the torsinA construct)/Percent of cells strongly exhibiting K48 foci in the controls)*100.

#### Fixed-cell epifluorescence microscopy

Fixed-cell epifluorescence imaging was performed either on a Nikon (Melville, NY) Eclipse Ni-E microscope or an Intelligent Imaging Innovations (Denver, CO) Marianas 200 Microscopy Workstation. The Ni-E microscope was driven by NIS-Elements software using a 40X /1.30 NA Plan Fluor oil immersion objective lens (Nikon Instruments Inc., Melville, NY), a SOLA solid state white-light excitation subsystem (Lumencor), and a CoolSNAP ES2 CCD camera (Photometrics). A custom DAPI filter set (#49028 with exciter: ET395/25x, dichroic: T425LPXR, and emitter: ET460/50m, Chroma Technology, Bellows Falls, VT) for the SOLA light source was used. EGFP (C-FL EGFP Zero Shift, #96362, Nikon Instruments Inc.), and Texas Red (C-FL Texas Red Zero Shift, #96365, Nikon Instruments Inc.) filter sets were also used. The Marianas 200 Microscopy Workstation was built on a Zeiss (Jena, Germany) AxioObserver Z.1 stand and driven by SlideBook 6.0 (Intelligent Imaging Innovations (3I), Denver, CO) using a 40x Plan-Apochromat oil immersion objective (NA=1.4, Zeiss) M27 with a DIC III Prism, a 300W Xenon 2-Galvo Lambda DG4 Fast Illuminator light source (Sutter Instrument, Novato, CA), a Photometrics CoolSnap HQ2 CCD (Tucson, AZ), and a BrightLine® Sedat filter set, and an optimized for DAPI, FITC, TRITC, and Cy5 (Semrock, Rochester, NY).

#### Brightness measurement protocol

U2OS cells were grown in 24-well glass bottom plates (In Vitro Scientific, Sunnyvale, CA) 12∼24 hours before transfection. Their media was then replaced with Dulbecco’s phosphate-buffered saline containing calcium and magnesium (BioWhittaker, Walkerville, MD) before measurement. Transiently transfected cells were selected using epifluorescence followed by a z-scan to ensure proper sub-cellular localization of the labeled protein (Hennen et al., 2017). The two-photon beam was then focused on the dorsal NE followed by the collection of intensity fluctuations for 1 minute. A second FFS measurement was then performed at the ventral NE. The brightness *λ_EGFP_* of the EGFP tag was determined from calibration measurements performed as previously described (Hennen et al., 2018a). Further details on the procedure for measuring and analyzing NE brightness can be found in Hennen et al. (2018) (Hennen et al., 2018a).

#### FFS experimental setup

FFS measurements were performed using a homebuilt two-photon microscope (Hennen et al., 2018a). 1000 nm light was focused through a 63x C-Apochromat water immersion objective with numerical aperture (NA) = 1.2 (Zeiss, Oberkochen, Germany), with an average excitation power of 0.3-0.4 mW after the objective. Emitted photon counts were collected on an avalanche photodiode (SPCM-AQ-141 APD; Perkin-Elmer, Dumberry, Quebec, Canada) and recorded with a sampling rate of 20 kHz using a Flex04-12D card (correlator.com, Bridgewater, NJ). The detected counts were analyzed using programs written in IDL 8.7 (Research Systems, Boulder, CO). Brightness measurements were analyzed using bootstrapped time-shifted mean-segmented Q (tsMSQ) analysis as previously described (Hennen et al., 2019), while bleaching experiments were analyzed as described below. Z-scans were performed on a PZ2000 piezo stage (ASI, Eugene, OR) which was moved axially with an arbitrary waveform generator (33522 A; Agilent Technologies, Santa Clara, CA) supplying a linear ramp function with peak-to-peak amplitude of 1.6 V, corresponding to 24.1 μm of axial travel, and period of 10 s for a speed of 4.82 μm/s.

#### Brightness analysis

FFS data were analyzed using the previously described tsMSQ method (Hennen et al., 2018b). The *Q* values recovered from fits of the tsMSQ curves were converted to normalized brightness *b* via the two equations 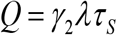 and 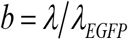, where *γ*_2_ is the PSF shape factor, *λ* is the brightness, and *τ_S_* is the photon count sampling time. Bias in the brightness caused by a decay in the intensity curves was estimated using a theoretical model that accounts for the depletion of fluorescence due to photobleaching (Hur and Mueller, 2015b). The characteristic intensity decay time and the decay amplitude are the critical parameters determining the level of bias. Using a typical decay time on the order of 10 s and the decay amplitudes observed for SS-EGFP-torsinA^WT^ (Fig. S2A) predicted a brightness bias of less than 10% for a data segment time of 6 s. Because the degree of bias decreases with shorter segment times (Hur and Mueller, 2015b), we separately calculated the brightness for data segment times of 1 s and 6 s. We obtained results that are identical within experimental uncertainty, which demonstrates the absence of bias. This analysis, which was performed for all brightness data presented in this study, found that a systematic error due to minor photobleaching was undetectable (Fig. S4).

#### Quantitative photobleaching measurements and analyses

The two-photon focus was centered onto the NE using reduced excitation power to avoid photobleaching while focusing. Photon counts were recorded upon exiting the low power mode. The fluorescence intensity decay was interpreted assuming a mobile and immobile population with fluorescence intensities of *F_m_* and *F_i_*(*t*), respectively. The mobile population diffuses sufficiently fast across the two-photon beam to make photobleaching negligible, which results in a time-independent and constant intensity *F_m_*. The immobile population with an initial intensity *F_i0_* = *F_i_*(0) decays over the course of the measurement to a final intensity of *F_i_*(∞) = 0, indicating complete photobleaching of its populations. Thus, the total fluorescence intensity initially contains the summed contributions of both populations, *F_0_* = *F_i0_*+ *F_m_*, and decays to a final amplitude of *F*_∞_ = *F_m_*. The fluorescence intensities *F_m_* = *λ_EGFP_N_m_* and *F_i_* = *λ_EGFP_N_i_* are converted into their respective number densities *N*_i_ and *N*_m_ by dividing by the brightness *λ_EGFP_* of the fluorescence label (Hennen et al., 2018c).

#### Polymer growth model

A simple polymer growth model for protein X assembling at an immobile seed site M is introduced. An initially mobile X diffusing along a nuclear membrane within the perinuclear space of the NE becomes immobilized upon binding to M, which is denoted by the species MX. A second X may bind to the immobilized MX, which leads to the species MX_2_. Further binding events increase the number of polymerized proteins X, with MX*_n_* denoting a complex of *n* immobile proteins at an immobile seed site, with *n* being a non-negative integer. The polymerization process is modeled using a simple equilibrium model with a single dissociation coefficient *K_D_*,

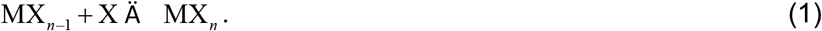

This equilibrium of this reaction is [MX*_n_*] = [M]([X]/*K_D_*)*^n^*, where the concentrations of M, X and MX*_n_* are denoted by [M], [X] and [MX*_n_*], respectively. The value of *n* determines the degree of polymerization. The probability that *n* proteins occupy a seed site is given by 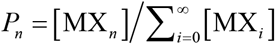, which determines the average degree of polymerization, 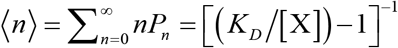. The next step is to rewrite this equation in terms of the average number of mobile and immobile proteins (*N_m_* and *N_i_*) found within the PSF area. The cross-sectional area A_PSF_ relates the concentration [X] to the number of mobile proteins within the PSF by *N_m_* = [X] A*_PSF_*. The number of immobile proteins within the PSF is related to the average degree of polymerization by where *S* is the number of seed sites per A_PSF_. We further express the dissociation constant in units of number

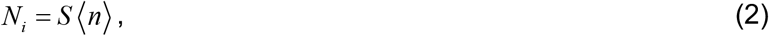

where *S* is the number of seed sites per A_PSF_. We further express the dissociation constant in units of number density, *K* = *K_D_A_PSF_* (Hennen et al., 2018c). With these definitions, the polymerization model is rewritten as

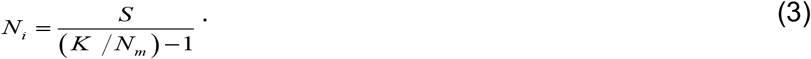

This binding model is mathematically equivalent to the uncommon type III gas adsorption and desorption isotherm (Khalfaoui et al., 2003). Eq. 3 is characterized by an asymptotic increase of the immobile population to infinity as the mobile population approaches *K*, thus requiring *N_m_* < *K*.

### KEY RESOURCES TABLE

| REAGENT OR RESOURCE | SOURCE | IDENTIFIER |
| --- | --- | --- |
| **Antibodies** |  |  |
| Mouse monoclonal anti-EGFP | Sigma-Aldrich | JL-8 |
| Rabbit monoclonal anti-K48-Ub | Millipore | AB_11213655 |
| Rabbit polyclonal anti-pericentrin | Covance | PRB-432C |
| Rat monoclonal anti-tyrosinated α-tubulin | European Collection of Animal Cultures | YL1/2 |
| **Bacterial and Virus Strains** |  |  |
| *E. coli* DH5α | ThermoFischer Scientific | 18265017 |
| *E. coli* *dam*<sup>-</sup>/*dcm*<sup>-</sup> | New England Biolabs | C2925 |
| **Chemicals, Peptides, and Recombinant Proteins** |  |  |
| DAPI | ThermoFisher Scientific | D1306 |
| LPA | Avanti Polar Lipids | #857130P |
| Phusion® High-fidelity DNA polymerase | New England Biolabs | M0530S |
| **Critical Commercial Assays** |  |  |
| GeneJet Plasmid Midiprep Kit | ThermoFisher Scientific | K0481 |
| GenJet™ *In Vitro* DNA Transfection Reagent | SignaGen Laboratories | SL100488 |
| QuikChange II Site-Directed Mutagenesis Kit | Aligent Technologies | 200523 |
| Wizard SV Gel and PCR Clean-Up System | Promega | A9281 |
| **Experimental Models: Cell Lines** |  |  |
| *Tor1A*<sup>-/-</sup> MEFs | (Saunders et al., 2017) | N/A |
| *TorA/B/2A/3A*-KO HeLa cells | (Laudermilch et al., 2016) | N/A |
| U2OS cells | ATCC | ATCC® HTB-96™ |
| **Oligonucleotides** |  |  |
| See Supplemental Table 1 |  |  |
| **Recombinant DNA** |  |  |
| EGFP-LAP1<sup>WT</sup> | (Saunders et al., 2017) | N/A |
| EGFP-LULL1<sup>WT</sup> | This work | N/A |
| SS-EGFP-torsinA<sup>WT</sup> | (Goodchild and Dauer, 2004) | N/A |
| SS-EGFP-torsinA<sup>K108A</sup> | (Goodchild and Dauer, 2004) | N/A |
| SS-EGFP-torsinA<sup>E171Q</sup> | (Goodchild and Dauer, 2004) | N/A |
| SS-EGFP-torsinA<sup>ΔE302/303</sup> | (Goodchild and Dauer, 2004) | N/A |
| SS-torsinA<sup>WT</sup>-EGFP | (Naismith et al., 2004) | N/A |
| SS-EGFP-torsinA<sup>G251D</sup> | This work | N/A |
| SS-EGFP-torsinA<sup>ΔNTD</sup> | This work | N/A |
| SS-EGFP-torsinA<sup>2xLeu</sup> | This work | N/A |
| SS-EGFP-torsinA<sup>C49Y</sup> | This work | N/A |
| SS-EGFP-torsinA<sup>ΔNTD, K108A</sup> | This work | N/A |
| SS-EGFP-torsinA<sup>ΔNTD, E171Q</sup> | This work | N/A |
| SS-EGFP-torsinA<sup>ΔNTD, ΔE</sup> | This work | N/A |
| SS-EGFP-torsinA<sup>ΔNTD, G251D</sup> | This work | N/A |
| SS-EGFP-torsinA<sup>21-49</sup> | This work | N/A |
| SS-EGFP-torsinA<sup>50-332</sup> | This work | N/A |
| pSOTA | This work | N/A |
| pSOTA.1 | This work | N/A |
| pSOTA.2 | This work | N/A |
| pSOTA.3 | This work | N/A |
| pSOTA.3* | This work | N/A |
| pSOTA<sub>Entry</sub> | This work | N/A |
| pSOTA<sub>Entry</sub>torsinA<sub>2</sub><sup>WT*</sup> | This work | N/A |
| pSOTA<sub>Entry</sub>torsinA<sub>3</sub><sup>WT*</sup> | This work | N/A |
| pSOTA<sub>Entry</sub>torsinA<sub>4</sub><sup>WT*</sup> | This work | N/A |
| pSOTA<sub>Entry</sub>torsinA<sub>5</sub><sup>WT*</sup> | This work | N/A |
| pSOTA<sub>Entry</sub>torsinA<sub>6</sub><sup>WT*</sup> | This work | N/A |
| SS-EGFP-SOTA<sub>2</sub> | This work | N/A |
| SS-EGFP-SOTA<sub>3</sub> | This work | N/A |
| SS-EGFP-SOTA<sub>4</sub> | This work | N/A |
| SS-EGFP-SOTA<sub>5</sub> | This work | N/A |
| SS-EGFP-SOTA<sub>6</sub> | This work | N/A |
| **Software and Algorithms** |  |  |
| Excel | Microsoft | N/A |
| IDL | Harris Geospatial | N/A |
| ImageJ | NIH | N/A |
| MatLab | Softworks | N/A |
| NIS-Elements software | Nikon | N/A |
| SlideBook | 3I | N/A |
| Word | Microsoft | N/A |

---

**Supplemental Figure 1. Plots of *b* vs. *N* with extended range for SS-EGFP-torsinA^50-332^, SS-EGFP-torsinA^ΔNTD^, SS-EGFP-torsinA^ΔNTD, K108A^, and SS-EGFP-torsinA^ΔNTD, ΔE^.** A-D) Plots of *b* vs. *N* within the NE of U2OS cells expressing the indicated constructs.

**Supplemental Figure 2. Fluorescence intensity traces of SS-EGFP-torsinA^NTD-2xLeu^, SS-EGFP-SOTA2, SS-EGFP-SOTA3, SS-EGFP-SOTA4, SS-EGFP-SOTA5, and SS-EGFP-torsinA^WT^ in the NE of U2OS cells.** Intensity traces of the indicated constructs measured within the NE of U2OS cells. Dark and light gray circles represent two intensity traces taken in different cells. The higher expressing cell is represented by the dark gray symbols. A decay in the fluorescence intensity indicates the presence of photobleaching.

**Supplemental Figure 3. Model of the tubular torsinA seed sites within the NE.** A-B) Side (A) and top-down (B) views of SS-EGFP-SOTA6 constructs forming a helical polymer at each tubular seed side. The nuclear membrane extrudes into the inner wall formed by the helical filament. C-D) Side (C) and top-down (D) views of SS-EGFP-SOTA3 constructs forming a helical polymer at each tubular seed side. Individual SS-EGFP-SOTA constructs are represented by differently colored spheres and the nuclear membrane is represented in grey.

**Supplemental Figure 4. Brightness ratio of SS-EGFP-torsinA^WT^ data as a function of *N*.** The brightness of selected data from Fig. 1B was recalculated for data segment times of 1 s and 6 s. Both methods recovered identical brightness values as demonstrated by the plot of the brightness ratio *b*_6s_ / *b*_1s_ as a function of *N*, demonstrating that photobleaching artifacts are negligible.

**Table S1:**
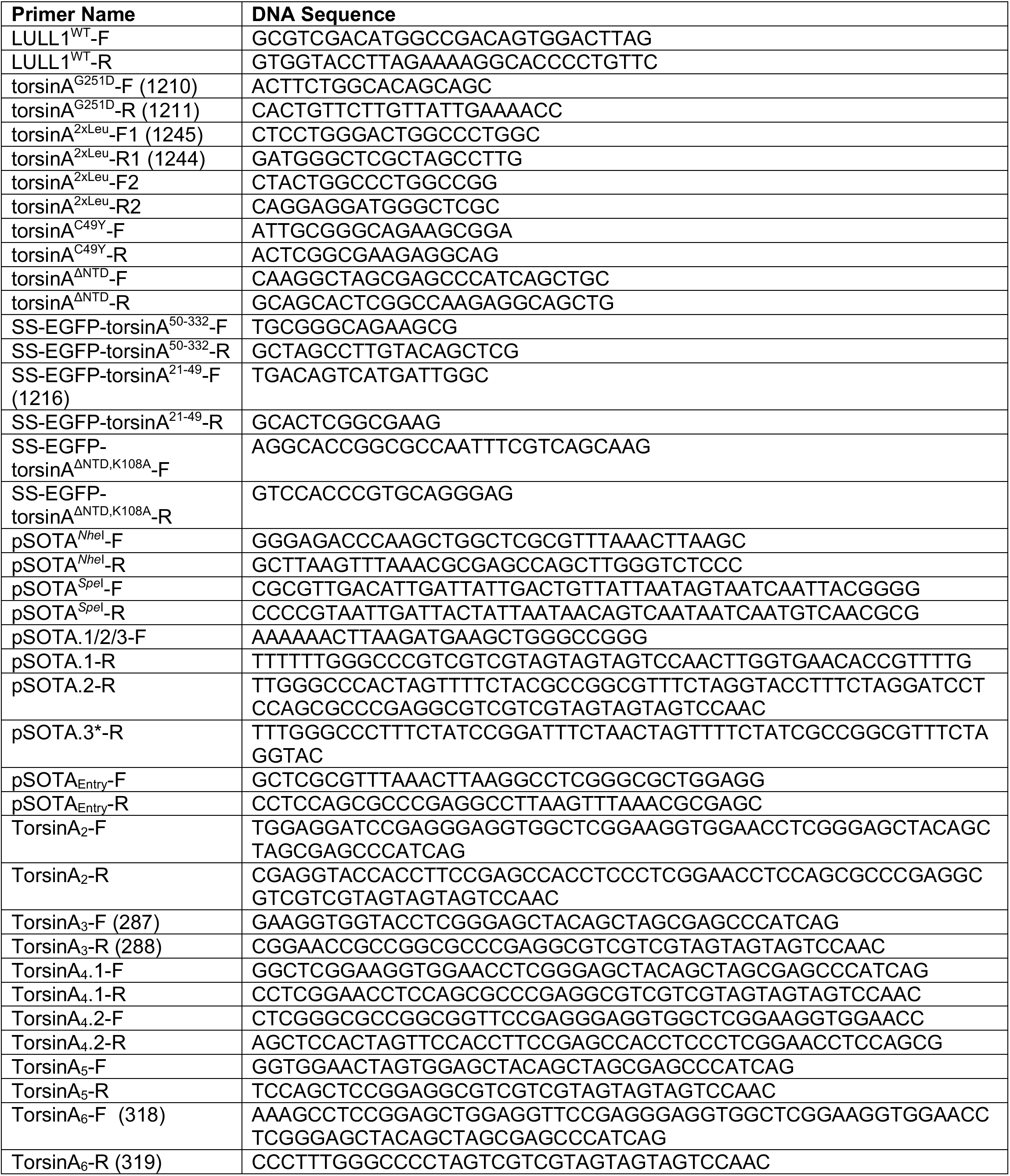
Primers used to generate the constructs used in this paper. The F or R in the primer name refers to forward or reverse, respectively. RE: restriction enzyme denoted by underlined sequence.

## REFERENCES

Bag, N., and Wohland, T. (2014) Imaging fluorescence fluctuation spectroscopy: new tools for quantitative bioimaging. Annu Rev Phys Chem, 65, 225–248.

Beauvais, G., Bode, N.M., Watson, J.L., Wen, H., Glenn, K.A., Kawano, H., Harata, N.C., Ehrlich, M.E., and Gonzalez-Alegre, P. (2016) Disruption of Protein Processing in the Endoplasmic Reticulum of DYT1 Knock-in Mice Implicates Novel Pathways in Dystonia Pathogenesis. J Neurosci, 36, 10245–10256.

Breakefield, X.O., Kamm, C., and Hanson, P.I. (2001) TorsinA: movement at many levels. Neuron, 31, 9–12.

Bressman, S.B., Sabatti, C., Raymond, D., de Leon, D., Klein, C., Kramer, P.L., Brin, M.F., Fahn, S., Breakefield, X., Ozelius, L.J. et al. (2000) The DYT1 phenotype and guidelines for diagnostic testing. Neurology, 54, 1746–1752.

Brown, R.S., Zhao, C., Chase, A.R., Wang, J., and Schlieker, C. (2014) The mechanism of Torsin ATPase activation. Proc Natl Acad Sci U S A, 111, E4822–31.

Burke, B., and Stewart, C.L. (2014) Functional architecture of the cell’s nucleus in development, aging, and disease. Curr Top Dev Biol, 109, 1–52.

Cain, N.E., Tapley, E.C., McDonald, K.L., Cain, B.M., and Starr, D.A. (2014) The SUN protein UNC-84 is required only in force-bearing cells to maintain nuclear envelope architecture. J Cell Biol, 206, 163–172.

Callan, A.C., Bunning, S., Jones, O.T., High, S., and Swanton, E. (2007) Biosynthesis of the dystonia-associated AAA+ ATPase torsinA at the endoplasmic reticulum. Biochem J, 401, 607–612.

Chalfant, M., Barber, K.W., Borah, S., Thaller, D., and Lusk, C.P. (2019) Expression of TorsinA in a heterologous yeast system reveals interactions with lumenal domains of LINC and nuclear pore complex components. Mol Biol Cell, 30, 530–541.

Chase, A.R., Laudermilch, E., and Schlieker, C. (2017a) Torsin ATPases: Harnessing Dynamic Instability for Function. Front Mol Biosci, 4, 29.

Chase, A.R., Laudermilch, E., Wang, J., Shigematsu, H., Yokoyama, T., and Schlieker, C. (2017b) Dynamic functional assembly of the Torsin AAA+ ATPase and its modulation by LAP1. Mol Biol Cell,

Chen, P., Burdette, A.J., Porter, J.C., Ricketts, J.C., Fox, S.A., Nery, F.C., Hewett, J.W., Berkowitz, L.A., Breakefield, X.O., Caldwell, K.A. et al. (2010) The early-onset torsion dystonia-associated protein, torsinA, is a homeostatic regulator of endoplasmic reticulum stress response. Hum Mol Genet, 19, 3502–3515.

Chen, Y., and Müller, J.D. (2007) Determining the stoichiometry of protein heterocomplexes in living cells with fluorescence fluctuation spectroscopy. Proc Natl Acad Sci U S A, 104, 3147–3152.

Chen, Y., Wei, L.N., and Müller, J.D. (2003) Probing protein oligomerization in living cells with fluorescence fluctuation spectroscopy. Proc Natl Acad Sci U S A, 100, 15492–15497.

Crisp, M., Liu, Q., Roux, K., Rattner, J.B., Shanahan, C., Burke, B., Stahl, P.D., and Hodzic, D. (2006) Coupling of the nucleus and cytoplasm: role of the LINC complex. J Cell Biol, 172, 41–53.

Cupo, R.R., and Shorter, J. (2020) Skd3 (human CLPB) is a potent mitochondrial protein disaggregase that is inactivated by 3-methylglutaconic aciduria-linked mutations. bioRxiv,

Demircioglu, F.E., Zheng, W., McQuown, A.J., Maier, N.K., Watson, N., Cheeseman, I.M., Denic, V., Egelman, E.H., and Schwartz, T.U. (2019) The AAA + ATPase TorsinA polymerizes into hollow helical tubes with 8.5 subunits per turn. Nat Commun, 10, 3262.

Dominguez Gonzalez, B., Billion, K., Rous, S., Pavie, B., Lange, C., and Goodchild, R. (2018) Excess LINC complexes impair brain morphogenesis in a mouse model of recessive TOR1A disease. Hum Mol Genet, 27, 2154–2170.

Esapa, C.T., Waite, A., Locke, M., Benson, M.A., Kraus, M., McIlhinney, R.A., Sillitoe, R.V., Beesley, P.W., and Blake, D.J. (2007) SGCE missense mutations that cause myoclonus-dystonia syndrome impair epsilon-sarcoglycan trafficking to the plasma membrane: modulation by ubiquitination and torsinA. Hum Mol Genet, 16, 327–342.

Fernandez-Martinez, J., and Rout, M.P. (2021) One Ring to Rule them All? Structural and Functional Diversity in the Nuclear Pore Complex. Trends Biochem Sci,

Gates, S.N., Yokom, A.L., Lin, J., Jackrel, M.E., Rizo, A.N., Kendsersky, N.M., Buell, C.E., Sweeny, E.A., Mack, K.L., Chuang, E. et al. (2017) Ratchet-like polypeptide translocation mechanism of the AAA+ disaggregase Hsp104. Science, 357, 273–279.

Gerace, L. (2004) TorsinA and torsion dystonia: Unraveling the architecture of the nuclear envelope. Proc Natl Acad Sci U S A, 101, 8839–8840.

Gonzalez-Alegre, P., and Paulson, H.L. (2004) Aberrant cellular behavior of mutant torsinA implicates nuclear envelope dysfunction in DYT1 dystonia. J Neurosci, 24, 2593–2601.

Goodchild, R.E., Buchwalter, A.L., Naismith, T.V., Holbrook, K., Billion, K., Dauer, W.T., Liang, C.C., Dear, M.L., and Hanson, P.I. (2015) Access of torsinA to the inner nuclear membrane is activity dependent and regulated in the endoplasmic reticulum. J Cell Sci, 128, 2854–2865.

Goodchild, R.E., and Dauer, W.T. (2004) Mislocalization to the nuclear envelope: an effect of the dystonia-causing torsinA mutation. Proc Natl Acad Sci U S A, 101, 847–852.

Goodchild, R.E., and Dauer, W.T. (2005) The AAA+ protein torsinA interacts with a conserved domain present in LAP1 and a novel ER protein. J Cell Biol, 168, 855–862.

Goodchild, R.E., Kim, C.E., and Dauer, W.T. (2005) Loss of the dystonia-associated protein torsinA selectively disrupts the neuronal nuclear envelope. Neuron, 48, 923–932.

Grillet, M., Dominguez Gonzalez, B., Sicart, A., Pöttler, M., Cascalho, A., Billion, K., Hernandez Diaz, S., Swerts, J., Naismith, T.V., Gounko, N.V. et al. (2016) Torsins Are Essential Regulators of Cellular Lipid Metabolism. Dev Cell, 38, 235–247.

György, B., Cruz, L., Yellen, D., Aufiero, M., Alland, I., Zhang, X., Ericsson, M., Fraefel, C., Li, Y.C., Takeda, S. et al. (2018) Mutant torsinA in the heterozygous DYT1 state compromises HSV propagation in infected neurons and fibroblasts. Sci Rep, 8, 2324.

Hanson, P.I., and Whiteheart, S.W. (2005) AAA+ proteins: have engine, will work. Nat Rev Mol Cell Biol, 6, 519–529.

Hennen, J., Angert, I., Hur, K.H., Gant Luxton, G.W., and Mueller, J.D. (2018a) Investigating LINC Complex Protein Homo-oligomerization in the Nuclear Envelopes of Living Cells Using Fluorescence Fluctuation Spectroscopy. Methods Mol Biol, 1840, 121–135.

Hennen, J., Hur, K.H., Karuka, S.R., Luxton, G.W.G., and Mueller, J.D. (2018b) Protein oligomerization and mobility within the nuclear envelope evaluated by the time-shifted mean-segmented Q factor. Methods,

Hennen, J., Hur, K.H., Kohler, J., Reddy Karuka, S., Angert, I., Luxton, G.W.G., and Mueller, J.D. (2020a) Identifying Heteroprotein Complexes in the Nuclear Envelope. Biophys J, 118, 26–35.

Hennen, J., Hur, K.H., and Mueller, J.D. (2019) Quantitative modeling of self-oligomerization of proteins in the nuclear envelope by fluorescence fluctuation analysis. Anal Biochem, 582, 113359.

Hennen, J., Hur, K.H., Saunders, C.A., Luxton, G.W.G., and Mueller, J.D. (2017) Quantitative Brightness Analysis of Protein Oligomerization in the Nuclear Envelope. Biophys J, 113, 138–147.

Hennen, J., Kohler, J., Karuka, S.R., Saunders, C.A., Luxton, G.W.G., and Mueller, J.D. (2020b) Differentiating Luminal and Membrane-Associated Nuclear Envelope Proteins. Biophys J,

Hennen, J., Saunders, C.A., Mueller, J.D., and Luxton, G.W.G. (2018c) Fluorescence Fluctuation Spectroscopy Reveals Differential SUN Protein Oligomerization In Living Cells. Mol Biol Cell,

Hewett, J., Ziefer, P., Bergeron, D., Naismith, T., Boston, H., Slater, D., Wilbur, J., Schuback, D., Kamm, C., Smith, N. et al. (2003) TorsinA in PC12 cells: localization in the endoplasmic reticulum and response to stress. J Neurosci Res, 72, 158–168.

Hewett, J.W., Kamm, C., Boston, H., Beauchamp, R., Naismith, T., Ozelius, L., Hanson, P.I., Breakefield, X.O., and Ramesh, V. (2004) TorsinB--perinuclear location and association with torsinA. J Neurochem, 89, 1186–1194.

Hewett, J.W., Nery, F.C., Niland, B., Ge, P., Tan, P., Hadwiger, P., Tannous, B.A., Sah, D.W., and Breakefield, X.O. (2008) siRNA knock-down of mutant torsinA restores processing through secretory pathway in DYT1 dystonia cells. Hum Mol Genet, 17, 1436–1445.

Hewett, J.W., Tannous, B., Niland, B.P., Nery, F.C., Zeng, J., Li, Y., and Breakefield, X.O. (2007) Mutant torsinA interferes with protein processing through the secretory pathway in DYT1 dystonia cells. Proc Natl Acad Sci U S A, 104, 7271–7276.

Hölper, J.E., Klupp, B.G., Luxton, G.W.G., Franzke, K., and Mettenleiter, T.C. (2020) Function of Torsin AAA+ ATPases in Pseudorabies Virus Nuclear Egress. Cells, 9,

Huelgas-Morales, G., Sanders, M., Mekonnen, G., Tsukamoto, T., and Greenstein, D. (2020) Decreased mechanotransduction prevents nuclear collapse in a Caenorhabditis elegans laminopathy. Proc Natl Acad Sci U S A, 117, 31301–31308.

Hur, K.H., Macdonald, P.J., Berk, S., Angert, C.I., Chen, Y., and Mueller, J.D. (2014) Quantitative measurement of brightness from living cells in the presence of photodepletion. PLoS One, 9, e97440.

Hur, K.H., and Mueller, J.D. (2015a) Quantitative Brightness Analysis of Fluorescence Intensity Fluctuations in E. Coli. PLoS One, 10, e0130063.

Hur, K.-H., and Mueller, J.D. (2015b) Quantitative brightness analysis of fluorescence intensity fluctuations in E. coli. PloS one, 10, e0130063.

Iqbal, Z., Koht, J., Pihlstrøm, L., Henriksen, S.P., Cappelletti, C., Russel, M.B., Norberto de Souza, O., Skogseid, I.M., and Toft, M. (2019) Missense mutations in DYT-TOR1A dystonia. Neurol Genet, 5, e343.

Jahed, Z., Soheilypour, M., Peyro, M., and Mofrad, M.R. (2016) The LINC and NPC relationship - it’s complicated. J Cell Sci, 129, 3219–3229.

Jokhi, V., Ashley, J., Nunnari, J., Noma, A., Ito, N., Wakabayashi-Ito, N., Moore, M.J., and Budnik, V. (2013) Torsin mediates primary envelopment of large ribonucleoprotein granules at the nuclear envelope. Cell Rep, 3, 988–995.

Jungwirth, M., Dear, M.L., Brown, P., Holbrook, K., and Goodchild, R. (2010) Relative tissue expression of homologous torsinB correlates with the neuronal specific importance of DYT1 dystonia-associated torsinA. Hum Mol Genet, 19, 888–900.

Jungwirth, M.T., Kumar, D., Jeong, D.Y., and Goodchild, R.E. (2011) The nuclear envelope localization of DYT1 dystonia torsinA-ΔE requires the SUN1 LINC complex component. BMC Cell Biol, 12, 24.

Kariminejad, A., Dahl-Halvarsson, M., Ravenscroft, G., Afroozan, F., Keshavarz, E., Goullée, H., Davis, M.R., Faraji Zonooz, M., Najmabadi, H., Laing, N.G. et al. (2017) TOR1A variants cause a severe arthrogryposis with developmental delay, strabismus and tremor. Brain, 140, 2851–2859.

Khalfaoui, M., Knani, S., Hachicha, M.A., and Lamine, A.B. (2003) New theoretical expressions for the five adsorption type isotherms classified by BET based on statistical physics treatment. Journal of colloid and interface science, 263, 350–356.

Kim, A.Y., Seo, J.B., Kim, W.T., Choi, H.J., Kim, S.Y., Morrow, G., Tanguay, R.M., Steller, H., and Koh, Y.H. (2015) The pathogenic human Torsin A in Drosophila activates the unfolded protein response and increases susceptibility to oxidative stress. BMC Genomics, 16, 338.

Kim, C.E., Perez, A., Perkins, G., Ellisman, M.H., and Dauer, W.T. (2010) A molecular mechanism underlying the neural-specific defect in torsinA mutant mice. Proc Natl Acad Sci U S A, 107, 9861–9866.

Kite, G.L. (1913) The relative permeability of the surface and interior portions of the cytoplasm of animal and plant cells. The Biological Bulletin, 25,

Knockenhauer, K.E., and Schwartz, T.U. (2016) The Nuclear Pore Complex as a Flexible and Dynamic Gate. Cell, 164, 1162–1171.

Laudermilch, E., Tsai, P.L., Graham, M., Turner, E., Zhao, C., and Schlieker, C. (2016) Dissecting Torsin/cofactor function at the nuclear envelope: a genetic study. Mol Biol Cell, 27, 3964–3971.

Li, P., and Noegel, A.A. (2015) Inner nuclear envelope protein SUN1 plays a prominent role in mammalian mRNA export. Nucleic Acids Res, 43, 9874–9888.

Liang, C.C., Tanabe, L.M., Jou, S., Chi, F., and Dauer, W.T. (2014) TorsinA hypofunction causes abnormal twisting movements and sensorimotor circuit neurodegeneration. J Clin Invest, 124, 3080–3092.

Liu, Q., Pante, N., Misteli, T., Elsagga, M., Crisp, M., Hodzic, D., Burke, B., and Roux, K.J. (2007) Functional association of Sun1 with nuclear pore complexes. J Cell Biol, 178, 785–798.

Liu, Z., Zolkiewska, A., and Zolkiewski, M. (2003) Characterization of human torsinA and its dystonia-associated mutant form. Biochemical Journal,

Luithle, N., de Bos, J.U., Hovius, R., Maslennikova, D., Lewis, R.T., Ungricht, R., Fierz, B., and Kutay, U. (2020) Torsin ATPases influence chromatin interaction of the Torsin regulator LAP1. Elife, 9,

Maimon, T., Elad, N., Dahan, I., and Medalia, O. (2012) The human nuclear pore complex as revealed by cryo-electron tomography. Structure, 20, 998–1006.

Maric, M., Haugo, A.C., Dauer, W., Johnson, D., and Roller, R.J. (2014) Nuclear envelope breakdown induced by herpes simplex virus type 1 involves the activity of viral fusion proteins. Virology, 460-461, 128–137.

Maric, M., Shao, J., Ryan, R.J., Wong, C.S., Gonzalez-Alegre, P., and Roller, R.J. (2011) A functional role for TorsinA in herpes simplex virus 1 nuclear egress. J Virol, 85, 9667–9679.

Martin, A., Baker, T.A., and Sauer, R.T. (2005) Rebuilt AAA + motors reveal operating principles for ATP-fuelled machines. Nature, 437, 1115–1120.

Mészáros, N., Cibulka, J., Mendiburo, M.J., Romanauska, A., Schneider, M., and Köhler, A. (2015) Nuclear pore basket proteins are tethered to the nuclear envelope and can regulate membrane curvature. Dev Cell, 33, 285–298.

Naismith, T.V., Dalal, S., and Hanson, P.I. (2009) Interaction of torsinA with its major binding partners is impaired by the dystonia-associated DeltaGAG deletion. J Biol Chem, 284, 27866–27874.

Naismith, T.V., Heuser, J.E., Breakefield, X.O., and Hanson, P.I. (2004) TorsinA in the nuclear envelope. Proc Natl Acad Sci U S A, 101, 7612–7617.

Nery, F.C., Armata, I.A., Farley, J.E., Cho, J.A., Yaqub, U., Chen, P., da Hora, C.C., Wang, Q., Tagaya, M., Klein, C. et al. (2011) TorsinA participates in endoplasmic reticulum-associated degradation. Nat Commun, 2, 393.

Nery, F.C., Zeng, J., Niland, B.P., Hewett, J., Farley, J., Irimia, D., Li, Y., Wiche, G., Sonnenberg, A., and Breakefield, X.O. (2008) TorsinA binds the KASH domain of nesprins and participates in linkage between nuclear envelope and cytoskeleton. J Cell Sci, 121, 3476–3486.

Otsuka, S., Bui, K.H., Schorb, M., Hossain, M.J., Politi, A.Z., Koch, B., Eltsov, M., Beck, M., and Ellenberg, J. (2016) Nuclear pore assembly proceeds by an inside-out extrusion of the nuclear envelope. Elife, 5,

Ozelius, L.J., Hewett, J.W., Page, C.E., Bressman, S.B., Kramer, P.L., Shalish, C., de Leon, D., Brin, M.F., Raymond, D., Corey, D.P. et al. (1997) The early-onset torsion dystonia gene (DYT1) encodes an ATP-binding protein. Nat Genet, 17, 40–48.

Pappas, S.S., Liang, C.C., Kim, S., Rivera, C.O., and Dauer, W.T. (2018) TorsinA dysfunction causes persistent neuronal nuclear pore defects. Hum Mol Genet, 27, 407–420.

Park, J., Damrauer, S.M., Baras, A., Reid, J.G., Overton, J.D., and Gonzalez-Alegre, P. (2019) Epidemiology of DYT1 dystonia: Estimating prevalence via genetic ascertainment. Neurol Genet, 5, e358.

Patel, S.S., and Rexach, M.F. (2008) Discovering novel interactions at the nuclear pore complex using bead halo: a rapid method for detecting molecular interactions of high and low affinity at equilibrium. Molecular & Cellular Proteomics, 7, 121–131.

Puchades, C., Sandate, C.R., and Lander, G.C. (2020) The molecular principles governing the activity and functional diversity of AAA+ proteins. Nat Rev Mol Cell Biol, 21, 43–58.

Rampello, A.J., Laudermilch, E., Vishnoi, N., Prophet, S.M., Shao, L., Zhao, C., Lusk, C.P., and Schlieker, C. (2020a) Torsin ATPase deficiency leads to defects in nuclear pore biogenesis and sequestration of MLF2. J Cell Biol, 219,

Rampello, A.J., Prophet, S.M., and Schlieker, C. (2020b) The Role of Torsin AAA+ Proteins in Preserving Nuclear Envelope Integrity and Safeguarding Against Disease. Biomolecules, 10,

Reichert, S.C., Gonzalez-Alegre, P., and Scharer, G.H. (2017) Biallelic TOR1A variants in an infant with severe arthrogryposis. Neurol Genet, 3, e154.

Saunders, C.A., Harris, N.J., Willey, P.T., Woolums, B.M., Wang, Y., McQuown, A.J., Schoenhofen, A., Worman, H.J., Dauer, W.T., Gundersen, G.G. et al. (2017) TorsinA controls TAN line assembly and the retrograde flow of dorsal perinuclear actin cables during rearward nuclear movement. J Cell Biol, 216, 657–674.

Saunders, C.A., and Luxton, G.W. (2016) LINCing defective nuclear-cytoskeletal coupling and DYT1 dystonia. Cell Mol Bioeng, 9, 207–216.

Shin, M., Puchades, C., Asmita, A., Puri, N., Adjei, E., Wiseman, R.L., Karzai, A.W., and Lander, G.C. (2020) Structural basis for distinct operational modes and protease activation in AAA+ protease Lon. Sci Adv, 6, eaba8404.

Siokas, V., Dardiotis, E., Tsironi, E.E., Tsivgoulis, G., Rikos, D., Sokratous, M., Koutsias, S., Paterakis, K., Deretzi, G., and Hadjigeorgiou, G.M. (2017) The Role of TOR1A Polymorphisms in Dystonia: A Systematic Review and Meta-Analysis. PLoS One, 12, e0169934.

Skinner, J.P., Chen, Y., and Müller, J.D. (2008) Fluorescence fluctuation spectroscopy in the presence of immobile fluorophores. Biophys J, 94, 2349–2360.

Slaughter, B.D., and Li, R. (2010) Toward quantitative “in vivo biochemistry” with fluorescence fluctuation spectroscopy. Mol Biol Cell, 21, 4306–4311.

Smith, E.M., Hennen, J., Chen, Y., and Mueller, J.D. (2015) In situ quantification of protein binding to the plasma membrane. Biophys J, 108, 2648–2657.

Sosa, B.A., Demircioglu, F.E., Chen, J.Z., Ingram, J., Ploegh, H.L., and Schwartz, T.U. (2014) How lamina-associated polypeptide 1 (LAP1) activates Torsin. Elife, 3, e03239.

Starr, D.A., and Fridolfsson, H.N. (2010) Interactions between nuclei and the cytoskeleton are mediated by SUN-KASH nuclear-envelope bridges. Annu Rev Cell Dev Biol, 26, 421–444.

Su, M., Guo, E.Z., Ding, X., Li, Y., Tarrasch, J.T., Brooks, C.L., Xu, Z., and Skiniotis, G. (2017) Mechanism of Vps4 hexamer function revealed by cryo-EM. Sci Adv, 3, e1700325.

Talamas, J.A., and Hetzer, M.W. (2011) POM121 and Sun1 play a role in early steps of interphase NPC assembly. J Cell Biol, 194, 27–37.

Tapley, E.C., Ly, N., and Starr, D.A. (2011) Multiple mechanisms actively target the SUN protein UNC-84 to the inner nuclear membrane. Mol Biol Cell, 22, 1739–1752.

Torres, G.E., Sweeney, A.L., Beaulieu, J.M., Shashidharan, P., and Caron, M.G. (2004) Effect of torsinA on membrane proteins reveals a loss of function and a dominant-negative phenotype of the dystonia-associated DeltaE-torsinA mutant. Proc Natl Acad Sci U S A, 101, 15650–15655.

Turner, E.M., Brown, R.S., Laudermilch, E., Tsai, P.L., and Schlieker, C. (2015) The Torsin Activator LULL1 Is Required for Efficient Growth of Herpes Simplex Virus 1. J Virol, 89, 8444–8452.

Ungricht, R., Klann, M., Horvath, P., and Kutay, U. (2015) Diffusion and retention are major determinants of protein targeting to the inner nuclear membrane. J Cell Biol, 209, 687–703.

Vander Heyden, A.B., Naismith, T.V., Snapp, E.L., and Hanson, P.I. (2011) Static retention of the lumenal monotopic membrane protein torsinA in the endoplasmic reticulum. EMBO J, 30, 3217–3231.

Vander Heyden, A.B., Naismith, T.V., Snapp, E.L., Hodzic, D., and Hanson, P.I. (2009) LULL1 retargets TorsinA to the nuclear envelope revealing an activity that is impaired by the DYT1 dystonia mutation. Mol Biol Cell, 20, 2661–2672.

VanGompel, M.J., Nguyen, K.C., Hall, D.H., Dauer, W.T., and Rose, L.S. (2015) A novel function for the Caenorhabditis elegans torsin OOC-5 in nucleoporin localization and nuclear import. Mol Biol Cell, 26, 1752–1763.

Vollmer, B., Lorenz, M., Moreno-Andrés, D., Bodenhöfer, M., De Magistris, P., Astrinidis, S.A., Schooley, A., Flötenmeyer, M., Leptihn, S., and Antonin, W. (2015) Nup153 recruits the Nup107-160 complex to the inner nuclear membrane for interphasic nuclear pore complex assembly. Developmental cell, 33, 717– 728.

Watson, M.L. (1955) The nuclear envelope: Its structure and relation to cytoplasmic membranes. The Journal of biophysical and biochemical cytology, 1, 257.

Watson, M.L. (1959) Further observations on the nuclear envelope of the animal cell. J Biophys Biochem Cytol, 6, 147–156.

Zehr, E., Szyk, A., Piszczek, G., Szczesna, E., Zuo, X., and Roll-Mecak, A. (2017) Katanin spiral and ring structures shed light on power stroke for microtubule severing. Nat Struct Mol Biol, 24, 717–725.

Zhao, C., Brown, R.S., Chase, A.R., Eisele, M.R., and Schlieker, C. (2013) Regulation of Torsin ATPases by LAP1 and LULL1. Proc Natl Acad Sci U S A, 110, E1545–54.

Zhao, C., Brown, R.S., Tang, C.H., Hu, C.C., and Schlieker, C. (2016) Site-specific Proteolysis Mobilizes TorsinA from the Membrane of the Endoplasmic Reticulum (ER) in Response to ER Stress and B Cell Stimulation. J Biol Chem, 291, 9469–9481.

Zhou, L., and Panté, N. (2010) The nucleoporin Nup153 maintains nuclear envelope architecture and is required for cell migration in tumor cells. FEBS Lett, 584, 3013–3020.

Zhu, L., Wrabl, J.O., Hayashi, A.P., Rose, L.S., and Thomas, P.J. (2008) The torsin-family AAA+ protein OOC-5 contains a critical disulfide adjacent to Sensor-II that couples redox state to nucleotide binding. Mol Biol Cell, 19, 3599–3612.

Zuleger, N., Kelly, D.A., Richardson, A.C., Kerr, A.R., Goldberg, M.W., Goryachev, A.B., and Schirmer, E.C. (2011) System analysis shows distinct mechanisms and common principles of nuclear envelope protein dynamics. J Cell Biol, 193, 109–123.

