## Supplementary figures and images for "The luminal AAA+ ATPase torsinA mediates distinct mechanisms of nuclear-cytoplasmic communication by adopting different functional assembly states"

### Supplemental Figures Hur, Hennen et al., (2021) bioRxiv

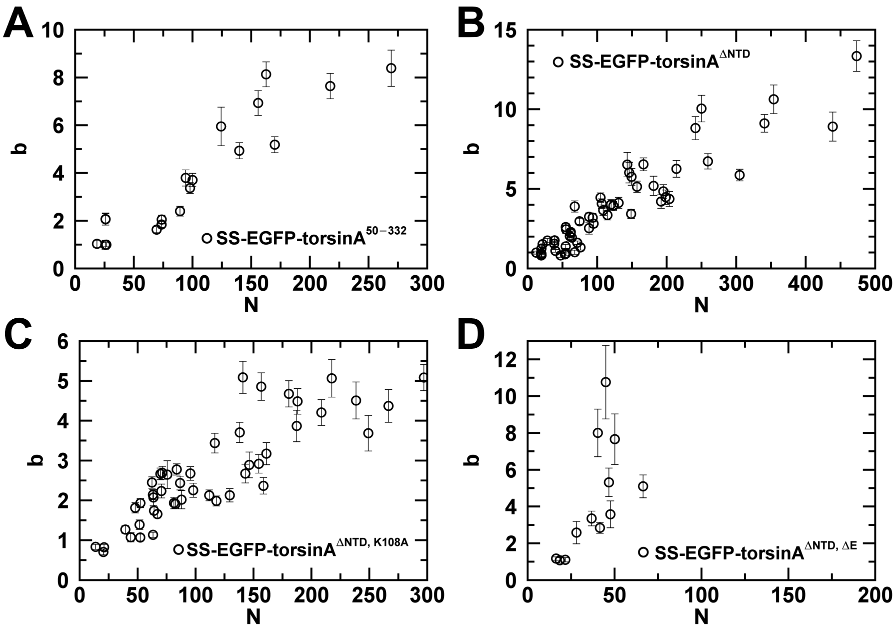

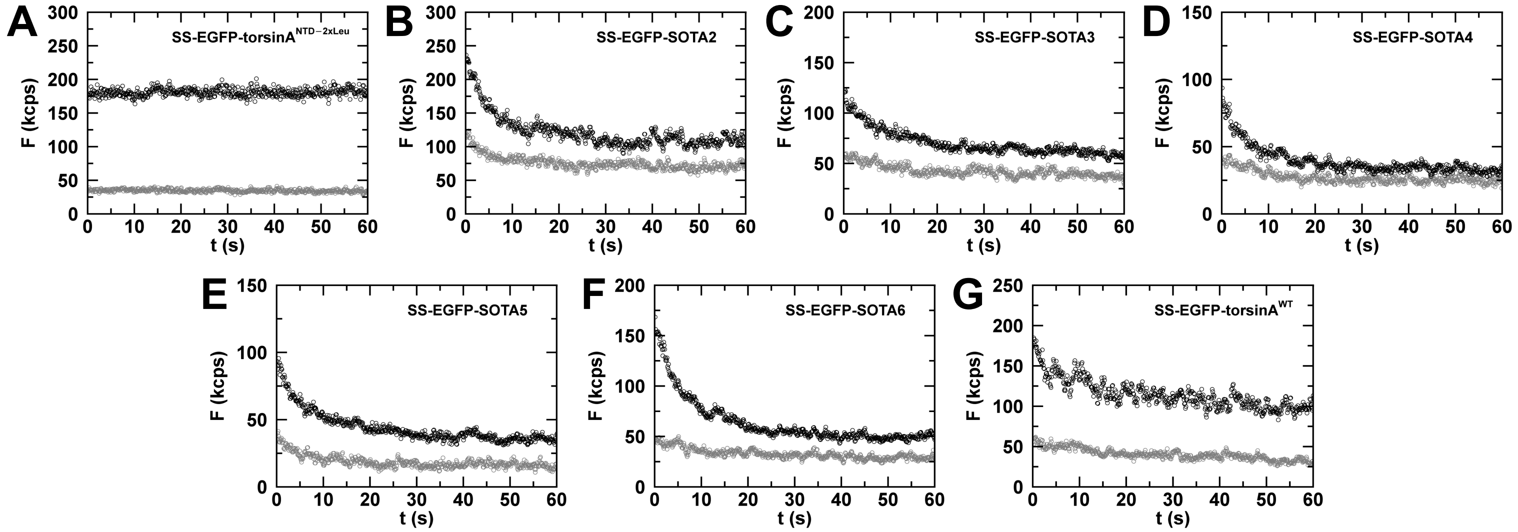

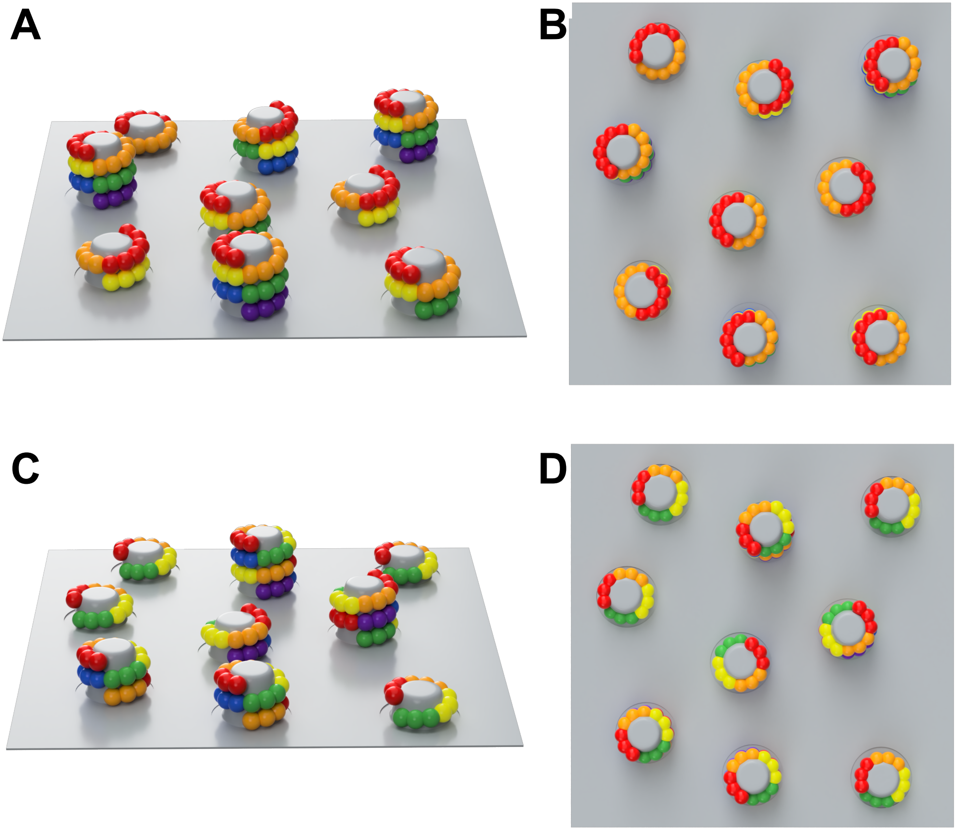


**Figure S3**

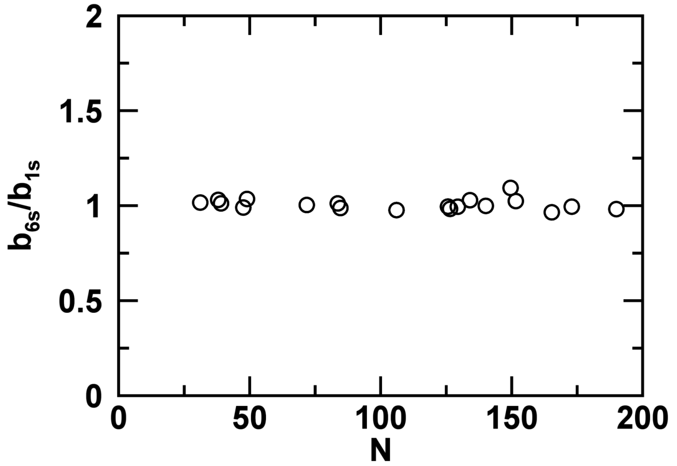


**Figure S4**
